# Statin-mediated reduction in mitochondrial cholesterol primes an anti-inflammatory response in macrophages by upregulating Jmjd3

**DOI:** 10.1101/2023.01.09.523264

**Authors:** Zeina Salloum, Kristin Dauner, Yun-fang Li, Neha Verma, John D. Zhang, Kiran Nakka, Mei Xi Chen, David Valdivieso-González, Víctor Almendro-Vedia, Jeffery McDonald, Chase D. Corley, Alexander Sorisky, Bao-Liang Song, Iván López-Montero, Jie Luo, Jeffrey F. Dilworth, Xiaohui Zha

## Abstract

Stains are known to be anti-inflammatory, but the mechanism remains poorly understood. Here we show that macrophages, either treated with statin *in vitro* or from statin-treated mice, have reduced cholesterol levels and higher expression of *Jmjd3,* a H3K27me3 demethylase. We provide evidence that lowering cholesterol levels in macrophages suppresses the ATP synthase in the inner mitochondrial membrane (IMM) and changes the proton gradient in the mitochondria. This activates NFκB and *Jmjd3* expression to remove the repressive marker H3K27me3. Accordingly, the epigenome is altered by the cholesterol reduction. When subsequently challenged by the inflammatory stimulus LPS (M1), both macrophages treated with statins *in vitro* or isolated from statin-treated mice *in vivo*, express lower levels pro-inflammatory cytokines than controls, while augmenting anti-inflammatory *Il10* expression. On the other hand, when macrophages are alternatively activated by IL4 (M2), statins promote the expression of *Arg1*, *Ym1*, and *Mrc1*. The enhanced expression is correlated with the statin-induced removal of H3K27me3 from these genes prior to activation. In addition, *Jmjd3* and its demethylase activity are necessary for cholesterol to modulate both M1 and M2 activation. We conclude that upregulation of *Jmjd3* is a key event for the anti-inflammatory function of statins on macrophages.

## Main Text

Many chronic diseases, including atherosclerosis, are associated with a low-grade level of inflammation (1). In atherosclerosis, macrophages are overloaded with cholesterol and are inflamed, which causes lesion formation (2). Statins, by inhibiting cholesterol biosynthesis and upregulating LDL receptor expression in hepatocytes and other cells, lower the level of circulating LDL. This leads to a decrease in the amount of cholesterol in peripheral tissues including macrophages (3–6). Statins reduce inflammation, but the mechanism for this effect is yet to be defined (7). Previously, we and others have observed that the level of cholesterol in resting macrophages, i.e., prior to stimulation, correlates directly with their pattern of inflammatory activation (8–10). For instance, when encountering lipopolysaccharides (LPS), cholesterol-rich macrophages release more inflammatory cytokines, such as TNF-α, IL-6, and IL12p40, and less anti-inflammatory cytokine IL-10, relative to control macrophages. Conversely, macrophages with reduced cholesterol content, such as those expressing ABCA1/G1 or being treated with statins, express fewer inflammatory cytokines, and more IL-10 upon identical LPS exposure (8–10). Noticeably, this association of cholesterol with specific type of macrophage inflammation is observed with multiple inflammatory stimuli, including ligands to TLR2, 3, 4 and other TRLs (9), implying a shared background in resting macrophages. It is known that, in order to mount a timely and vigorous defence against pathogens, macrophages activate several hundred genes immediately after sensing danger signals (11). This is achieved by employing a few select signal-dependent transcription factors, such as NF-κB, on a genome that is largely poised, i.e., epigenetically configured, prior stimulation (12). prior the stimulation (12). The organization of the epigenome in macrophages is initially defined by lineage-determining transcription factors and subsequently by metabolic /environmental cues, including those experienced in the past (12, 13). This forms a poised state in resting macrophages and allows the expression of context-dependent inflammatory genes (14, 15). We therefore speculate that cholesterol levels may directly influence the epigenome in resting macrophages, thereby influencing the inflammatory phenotype upon activation.

## Results

### Reducing cholesterol levels in macrophages activates the NF-κB pathway and upregulates Jmjd3, a histone demethylase

To test whether cholesterol levels alone can produce factors that change the epigenetic configuration in macrophages, we first treated RAW macrophages with statins^1^ for 2 days, which reduces cellular cholesterol by about 30% (8, 16). We also used 1 h methyl-β-cyclodextrin (MCD) treatment to acutely reduce cholesterol by a similar amount (8). This was to verify the cholesterol specificity of statins’ effects and also to provide a 1 h-protocol for inhibitor studies. RNA-seq was performed at the end of statin- or MCD-based treatments. As shown in Figure 1A, macrophages treated with statins altered the expression of a large number of genes (log2>1) (Figure 1A, a; Supplementary file 1; the top 40 upregulated genes: Figure 1-figure supplement 1, A). We applied the Gene Set Enrichment Analysis (GSEA) to all differentially expressed genes (up-and down-regulation), using the Hallmark genes database (Figure 1A, *b*). Genes of the NF-κB pathway activated by TNFα were the most highly represented group upon statin treatment (Figure 1A, b-c). Similarly, MCD-treated macrophages showed an identical highly represented group, i.e., NF-κB pathway activated by TNFα (Figure 1B, a-c; Supplementary file 2). The top 40 upregulated genes are in the supplementary figure (Figure 1-figure supplement 1, B). This analysis suggests that cholesterol depletion induces activation of NF-κB pathways in macrophages. Additional analysis, using a mouse transcriptional regulatory network database (17, 18), also identified the *Nfkb1* as the top TF in the promoters of genes upregulated by statins or MCD treatment (Figure 1-figure supplement 2, A & B). Genes down-regulated by statins, on another hand, are less enriched in TFs (Figure 1-figure supplement 2, C). In fact, MCD treatment produced no enrichment of TFs among down-regulated genes. Together, the RNA-sequencing data suggest that reducing cholesterol in macrophages primarily activates NF-κB pathways to enhance gene expression.

**Figure 1.**
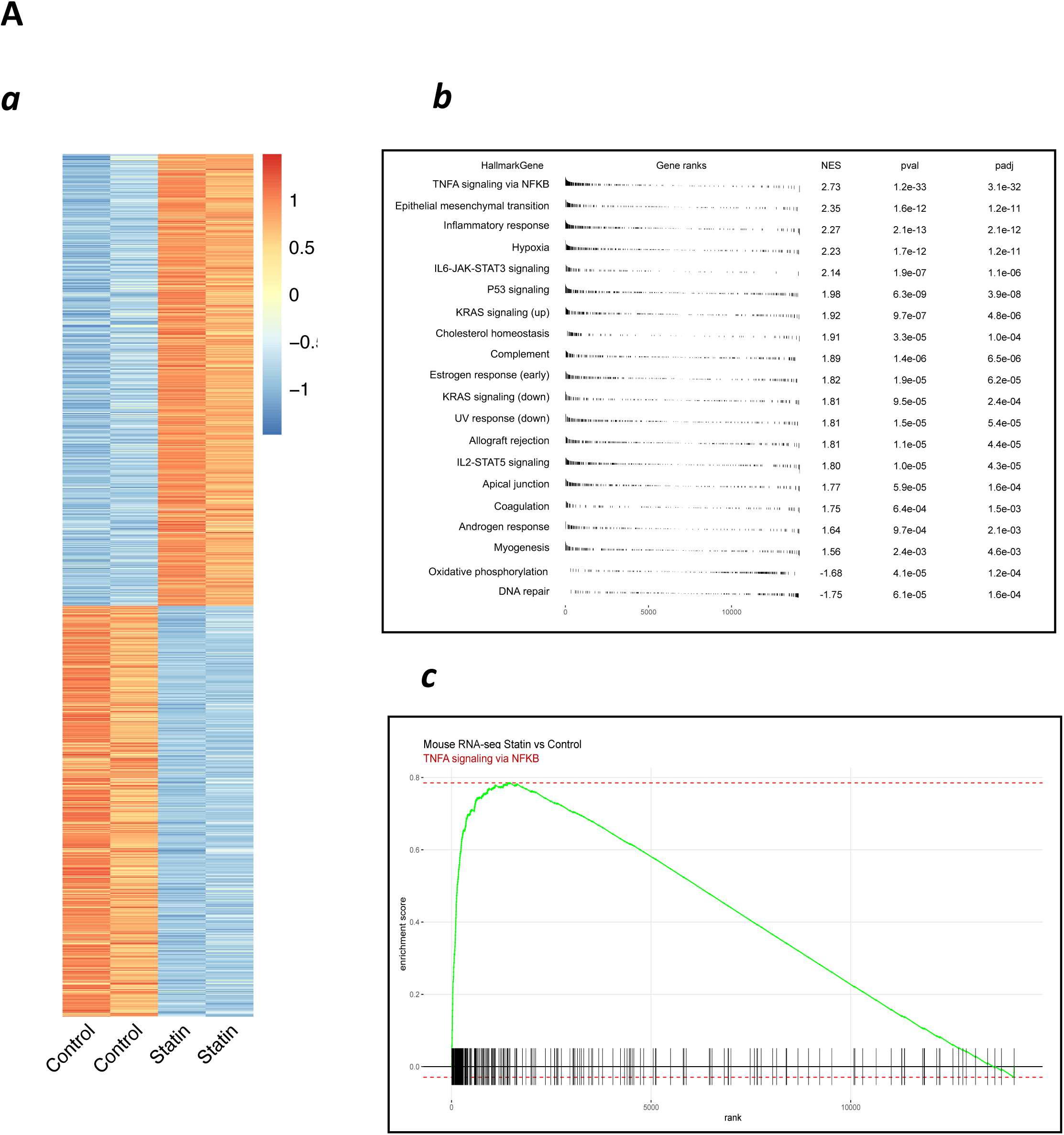
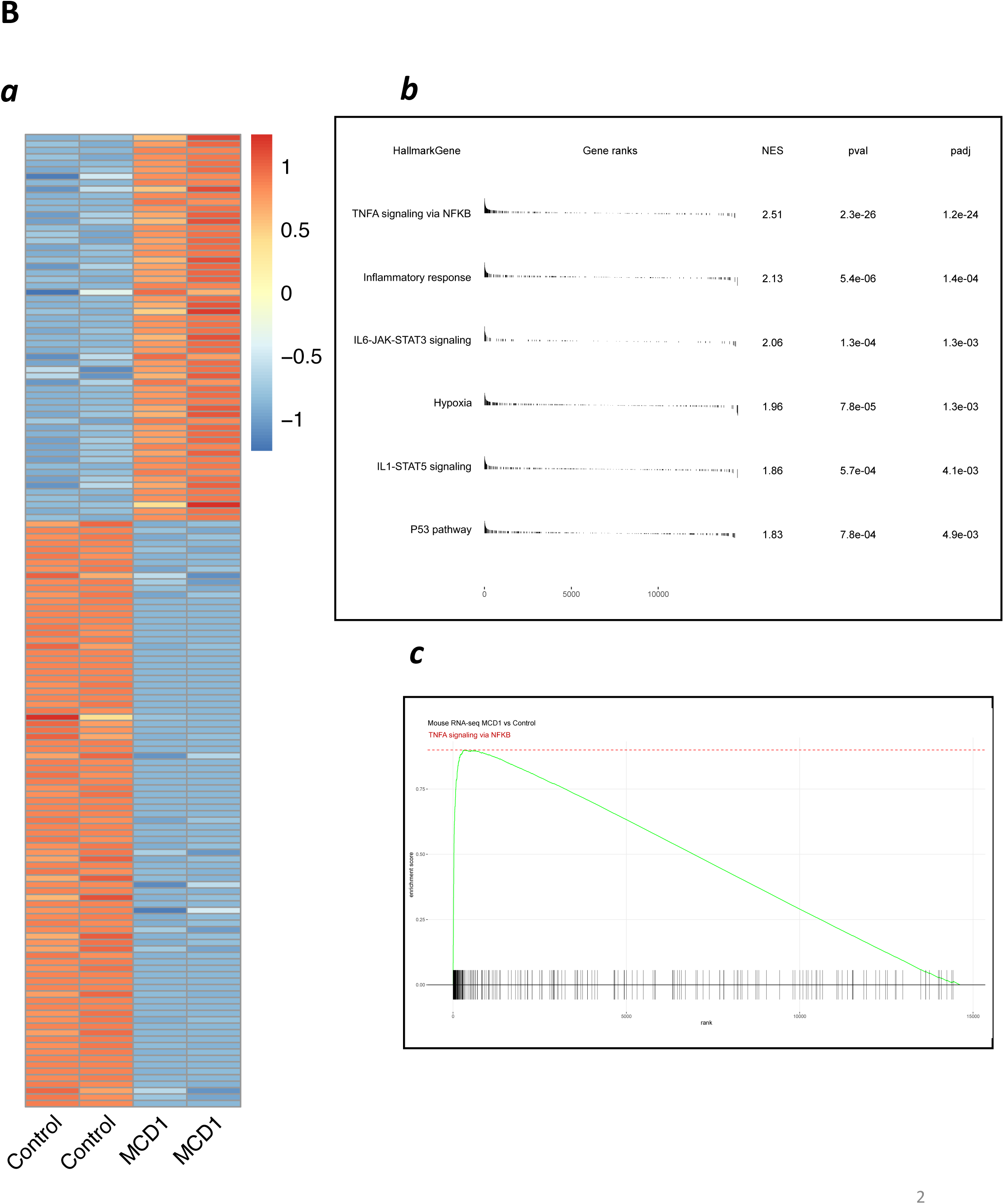
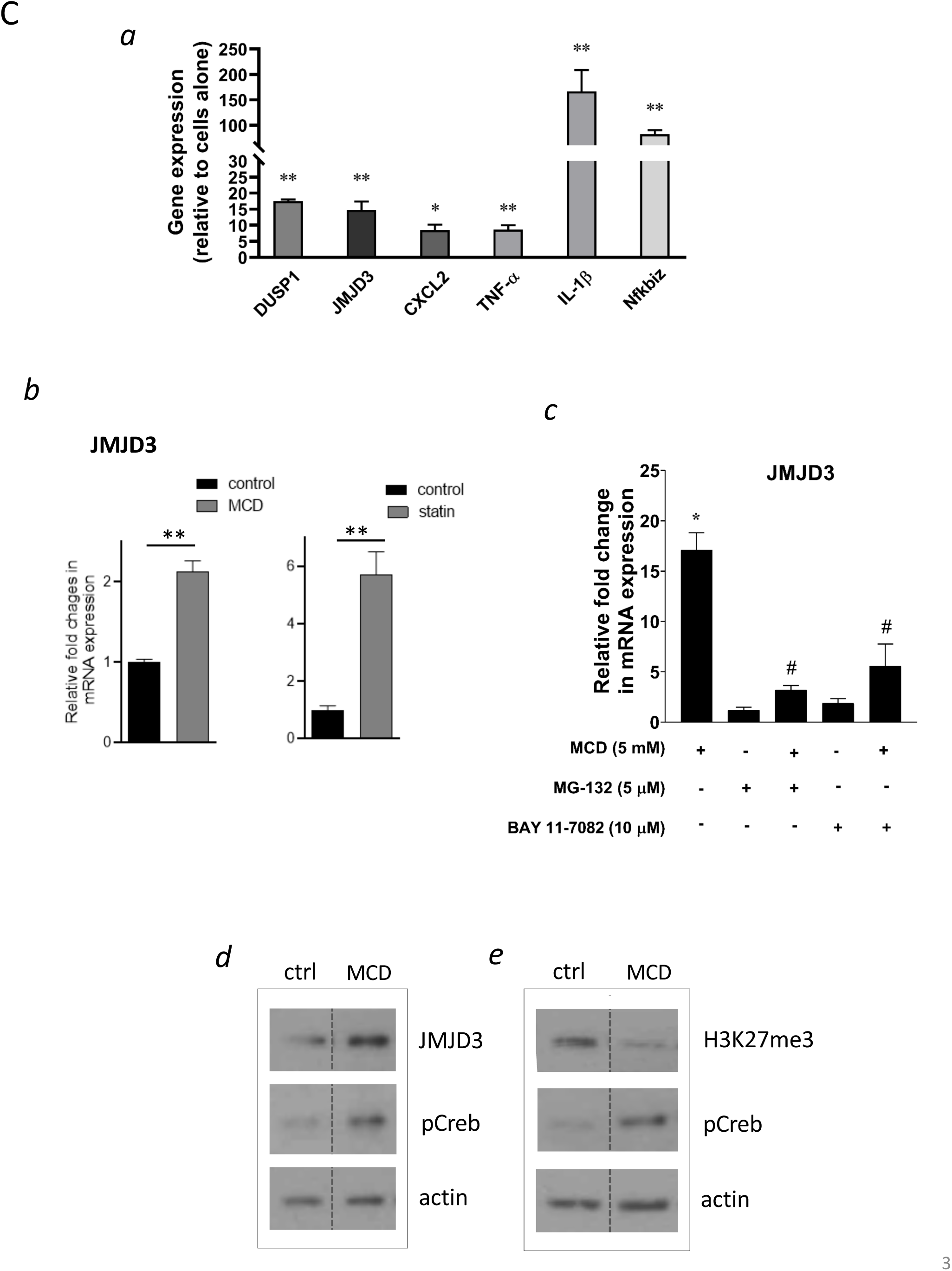
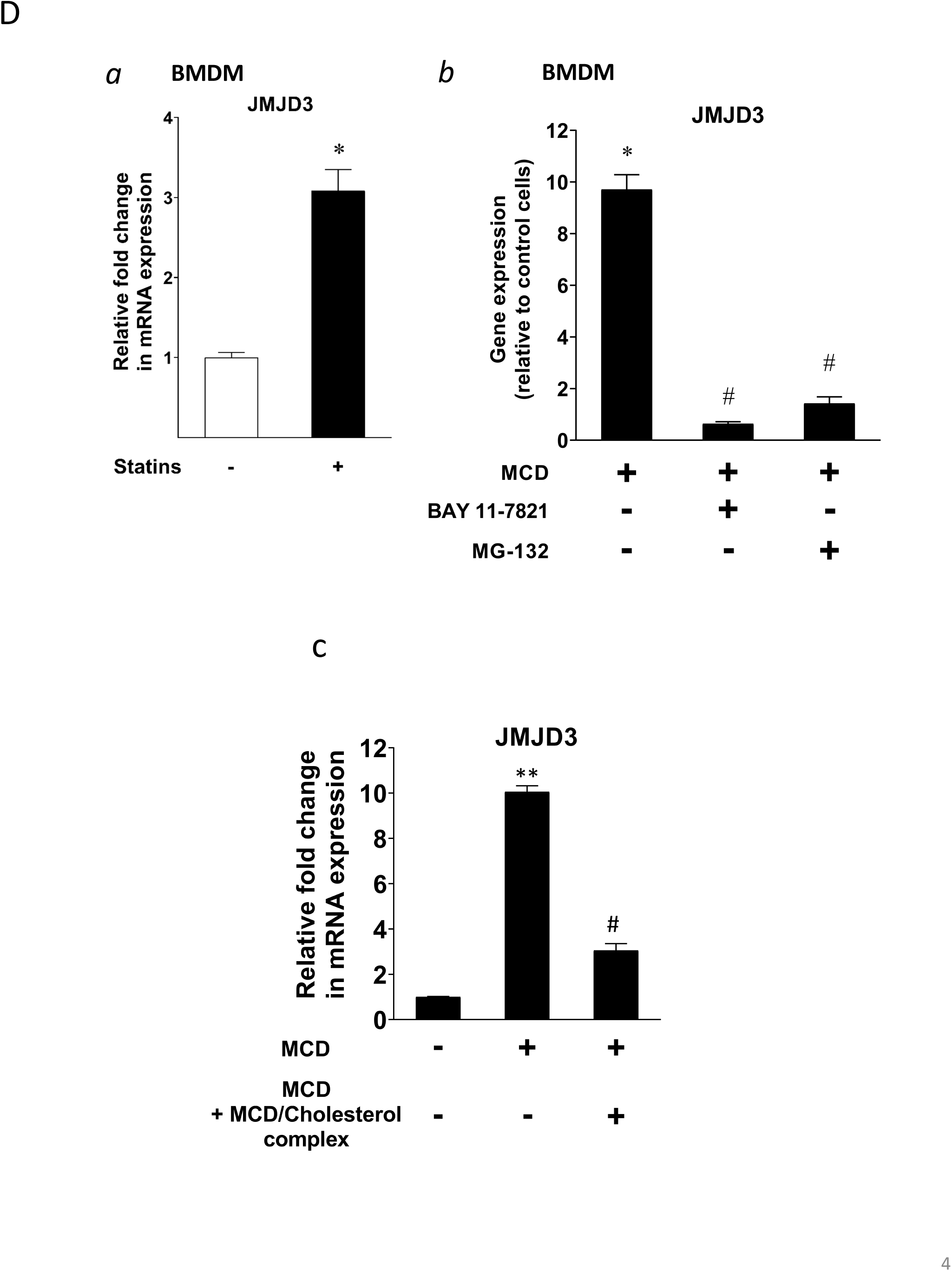
Satins upregulates the expression of *Jmjd3* in macrophages through NF-κB. (*A*) Statins activate NF-kB pathways in RAW 264.7 cells. *(a)* Heatmap of differentially expressed genes with or without statin (lovastatin, 7 µM + 200 µM mevalonate; 2 days). *(b)* Pathways identified by GSEA of differentially expressed genes in (*a*). *(c)* The details of most highly represented pathway, TNFA signaling vis NFKB. **(*B*)** MCD activate NF-kB pathways in RAW 264.7 cells. *(a)* Heatmap of differentially expressed genes with or without MCD (5 mM, 1 h). *(b)* Pathways identified by GSEA analysis of differentially expressed genes in (*a*). *(c)* The details of most highly represented pathway, TNFA signaling vis NFKB. **(*C*)** Statins and MCD upregulates *Jmjd3* in RAW 264.7 cells. *(a)* RT-qPCR of genes with or without MCD (5 mM; 1 h). *(b) Jmjd3* gene expression in MCD-, or statins-treated RAW 264.7 macrophages; *(**c**)* Effect of NF-kB inhibitors, MG-132 (5 µM) and BAY11-7082 (10 µM) on *Jmjd3* expression in MCD-treated RAW macrophages. *(d)* Western blotting of JMJD3 protein expression and *(e)* levels of H3K27Me3 in macrophages treated with 5 mM MCD (1h). The pCREB was used as internal control for cholesterol depletion and actin a loading control. Original blots are in source data. **(*D*)** Statin/MCD upregulates *Jmjd3* in BMDMs. *(a) Jmjd3* gene expression in statin-treated BMDMs (10 µM pravastatin + 200 µM mevalonate; 2 days); *(b*) Effect of NF-kB inhibitors, MG-132 [5 µM] and BAY11-7082 [10 µM], on *Jmjd3* expression in MCD-treated BMDMs. *(c) Jmjd3* expression in cholesterol repletes MCD-treated macrophages. Graphs are representative of 3 independent experiments with 3 replicates per condition and are presented as means ± SD. Statistical analysis was performed using unpaired, two-tailed Student’s t-test. An asterisk (*) or double asterisks (**) indicate a significant difference with p<0.05 and p<0.001, respectively. A hashtag (#) indicates a significant difference between MCD without or with inhibitors, p < 0.05.

We investigated several known NF-κB target genes using qPCR and noted that *Kdm6b*, encoding the H3K27me3 demethylase *Jmjd3* (19), was among the upregulated genes (Figure 1C, *a*). Both statins and MCD enhanced *Jmjd3* expression (Figure 1C, *b*). This upregulation was abolished by several structurally unrelated NF-κB inhibitors (Figure 1C, c and Figure 1-figure supplement 3), confirming that *Jmjd3* is a NF-κB target (20). We further confirmed that JMJD3 protein level is increased (Figure 1C, *d*) and that the level of H3K27me3, the substrate of JMJD3, is reciprocally decreased in macrophages with reduced cholesterol by MCD (Figure 1C, *e*). The elevated level of phosphorylated Creb (pCreb) is an indicator of effective cholesterol reduction (8). Similar results were seen in macrophages treated with statins (Figure 1-figure supplement 4). In addition, mouse bone marrow-derived macrophages (BMDMs) responded to cholesterol reduction identically: *Jmjd3* is upregulated by statins or MCD in a NF-κB-dependent manner (Figure 1D, *a & b*). Lastly, to verify the specificity of cholesterol in *Jmjd3* upregulation, MCD/cholesterol complex was used to replenish cellular cholesterol in MCD-treated BMDMs. This reversed the *Jmjd3* upregulation by cholesterol reduction in BMDMs (Figure 1D, c). We conclude that statin/MCD upregulates *Jmjd3* in macrophages likely through cholesterol reduction and NF-κB activation.

### Reducing cholesterol in macrophages directly activates NF-κB

GSEA above identifies the NF-κB pathway as the top activated biological process in both statin and MCD treated macrophages (Figure 1A & B). We thus directly examined NF-κB activities in macrophages treated with statin or MCD. Using RAW-Blue^TM^ cells containing a NF-κB reporter (21), we observed elevated NF-κB activity when cells were treated with MCD or statins (Figure 2A, *a & b*). In addition, two known NF-κB target genes, *ll1b* and *Tnfa*, were upregulated by MCD and statins, respectively (Figure 2A, *c & d*). Furthermore, blocking NF-κB function by inhibitors prevented the upregulation of *Il1b* and *Tnfa* by MCD (Figure 2A, *e & f* ). Lastly, to understand the scope of NF-κB activation by statins or MCD, we directly compared the magnitude of *Il1b* and *Tnfa* expression by statin or MCD with those by LPS. The effect of statins or MCD is much weaker than LPS: the changes in *Il1b* and *Tnfa* level of expression by statin/MCD are less than 1% of those stimulated by LPS (Figure 2B, *a & b*). We conclude that cholesterol reduction in macrophages likely activates NF-κB. However, this activation of NF-κB is of low magnitude, and distinct from the more robust NF-κB activation stimulated by LPS.

**Figure 2.**
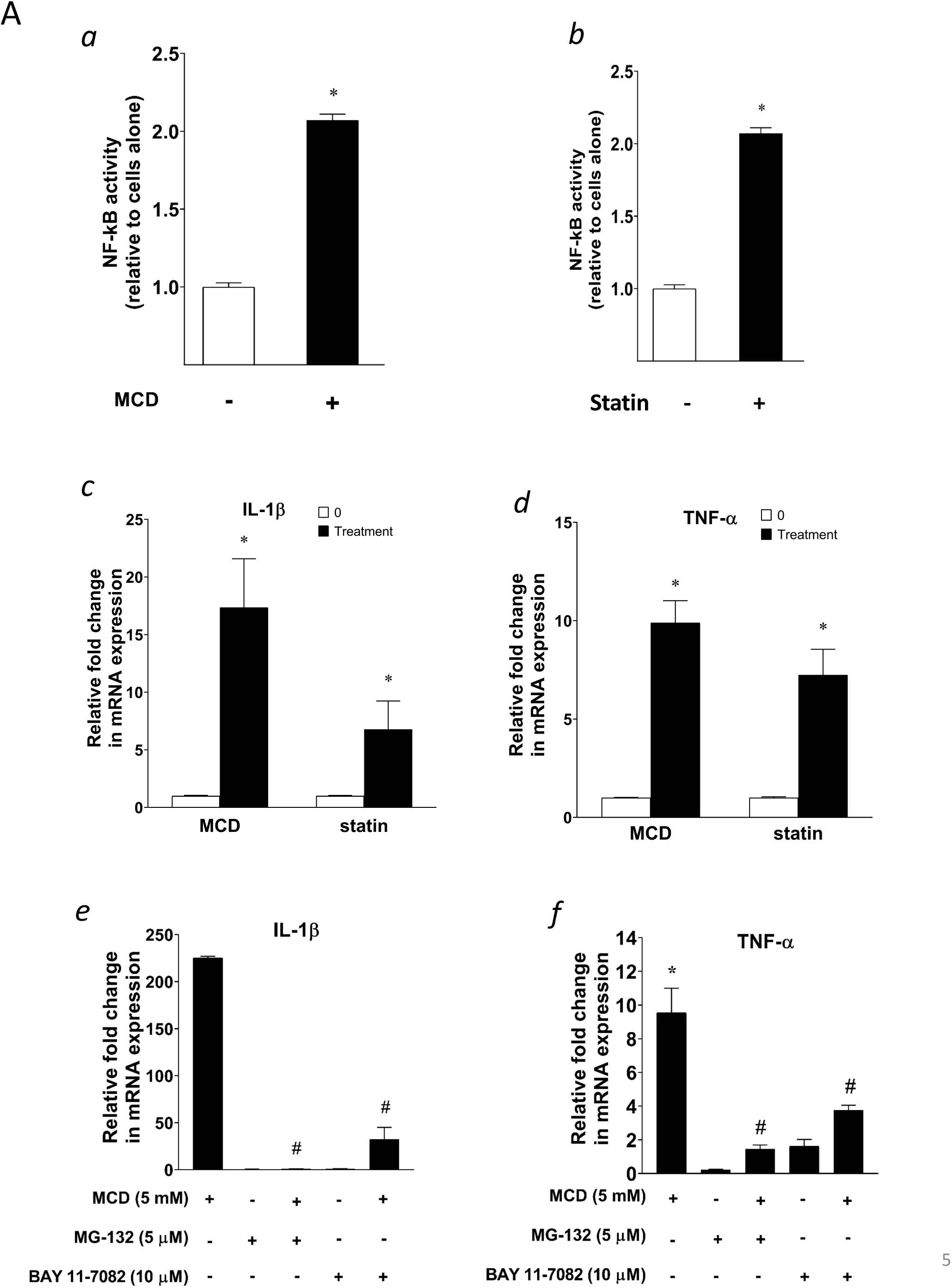
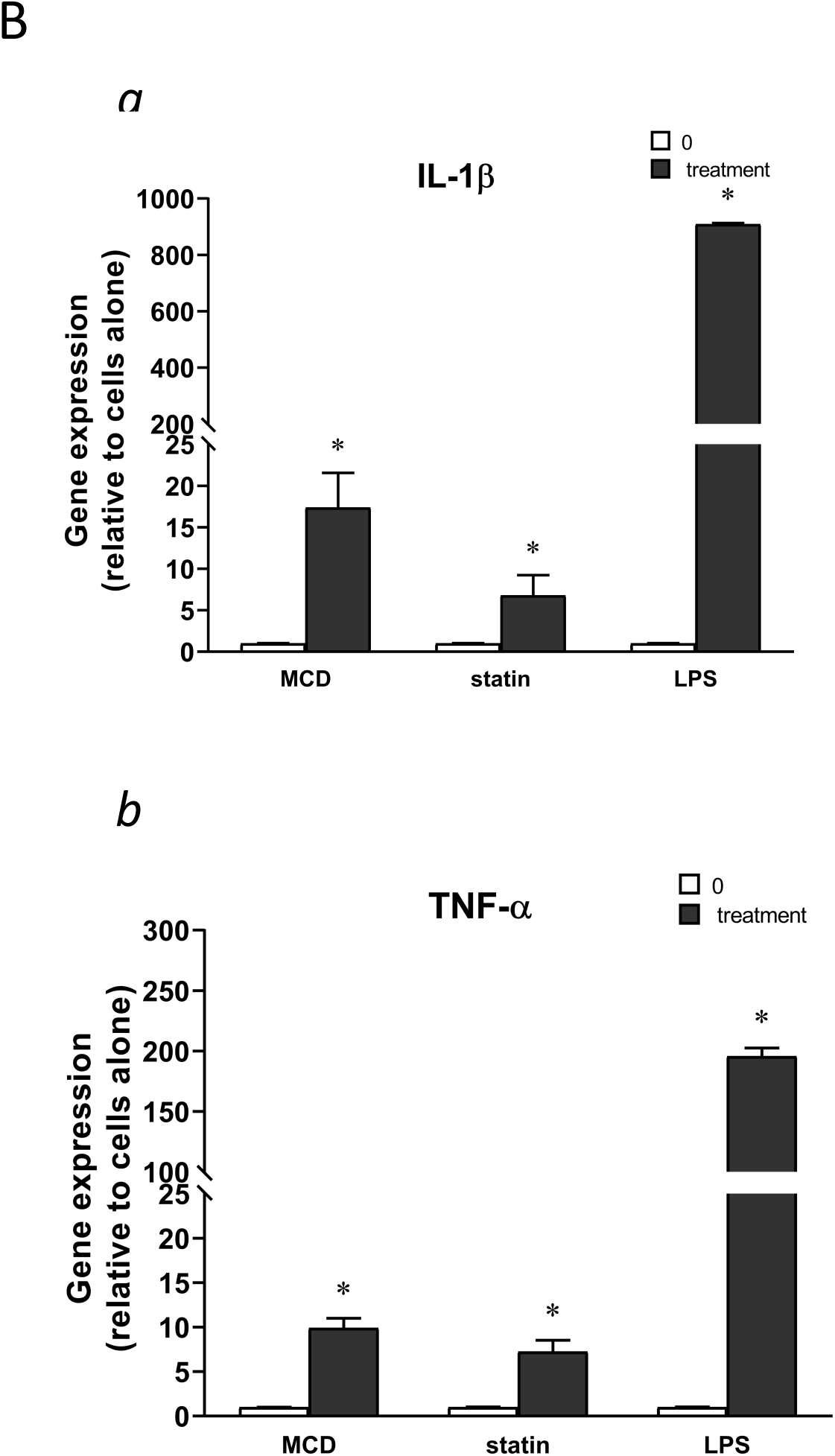
Cellular cholesterol contents regulate NF-kB pathway. (*A*) NF-kB activation by MCD *(a)* and statins *(b)* using RAW blue^TM^ macrophages. RT-qPCR of *Il1b (c)* and *Tnfa (d)* in RAW 264.7 cells with or without MCD (5 mM; 1 h), or with or without (10 µM compactin + 200 µM mevalonate; 2 days). Effect of NF-kB inhibitors (MG-132 [5 µM] and BAY11-7082 [10 µM]) on *Il1b (e)* and *Tnfa (f)* expression in MCD-treated RAW 264.7 macrophages. ***(B)*** The gene expression activated by MCD, statin or LPS. RT-qPCR of *Il1b (a)* and *Tnfa (b)* in RAW 264.7 cells with or without MCD (5 mM; 1 h), simvastatin (10 µM + 200 µM mevalonate; 2 days) or LPS (100 ng/ml; 3 h). Graphs are representative of 3 independent experiments with 3 replicates per condition and are presented as means ± SD. Statistical analysis was performed using unpaired, two-tailed Student’s t-test. An asterisk (*) or double asterisks (**) indicate a significant difference with p<0.05 and p<0.001, respectively. A hashtag (#) indicates a significant difference between MCD without or with inhibitors with p < 0.05.

### Reducing cholesterol in macrophages decreases mitochondria respiration

We next explored potential mechanisms by which cholesterol reduction with statin/MCD activates NF-κB. In recent years, it has become evident that NF-κB can be activated by a metabolic shift in the cell from oxidative phosphorylation (OXPHOS) in the mitochondria to glycolysis in the cytoplasm (22, 23). We therefore investigated whether statin/MCD modulates OXPHOS in macrophage to activate NF-κB. Using the extracellular flux analyzer (Seahorse), we found that 1 h MCD treatment of BMDMs decreased overall resting mitochondrial oxygen consumption rate (OCR) (Figure 3A, *a*). Moreover, this decrease was entirely attributable to the suppression of the ATP synthase (Figure 3A, *b*), a protein embedded in the inner mitochondrial membrane (IMM) and a component of electron transport chain. Interestingly, the maximal respiration remains unchanged by MCD (Figure 3A, *c*). Statin treatment similarly decreased overall resting mitochondrial OCR (Figure 3B, *a*) and that of ATP synthase (Figure 3B, *b*). However, statin also lowered the maximal respiration (Figure 3B, *c*), possibly due to the 2-day treatment period required for the statin treatment. Overall, statin or MCD suppresses ATP synthase in macrophages.

**Figure 3.**
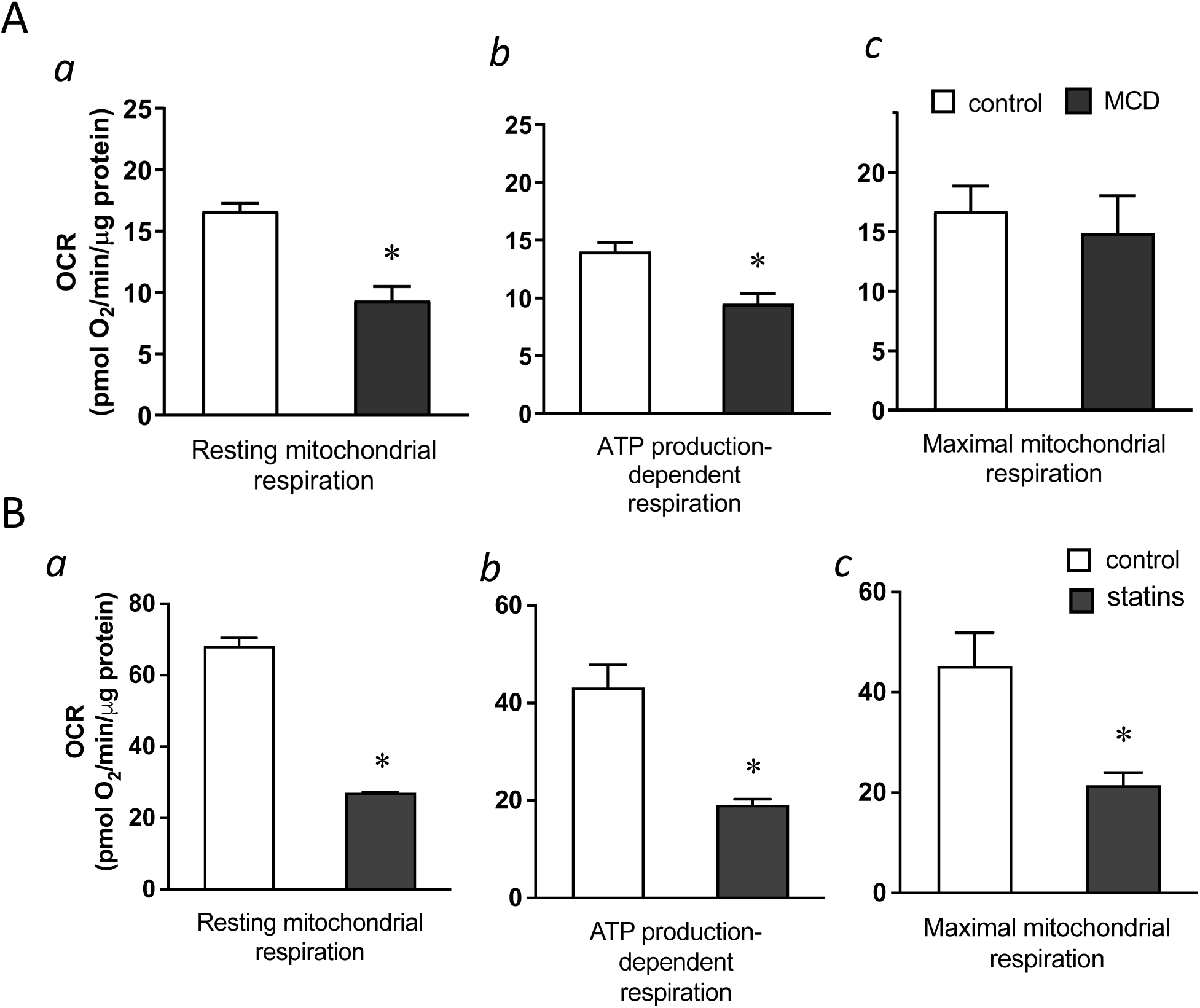
Cholesterol levels modulate mitochondrial respiration. (***A***) Mitochondrial oxygen consumption rates (OCR) of BMDMs treated with 1 h MCD (5 mM) or without. *(a)* OCR for mitochondrial resting respiration; *(b)* OCR representing mitochondrial ATP production-linked respiration. *(c)* OCR representing maximal respiration. ***(B)*** Mitochondrial oxygen consumption rates (OCR) in BMDMs treated compactin or without compactin (10 µM + 200 µM mevalonate; 2 days). *(a)* OCR for mitochondrial resting respiration; *(b)* OCR representing mitochondrial ATP production-linked respiration; *(c)* OCR representing maximal respiration. Data are representative of 3 independent experiments with 3 samples per group and data are presented as mean ± SD. Statistical analysis was performed using unpaired, two-tailed Student’s t-test. An asterisk (*) and (**) indicate a significant difference, p<0.05 and p<0.001.

### Reducing cholesterol in macrophages results in lower cholesterol level in IMM, which suppresses ATP synthase activity

We next asked how cholesterol reduction by statin/MCD might influence the ATP synthase in the IMM. In mammalian cells, the majority of cholesterol is in the plasma membrane. However, all intracellular membranes, including those in the mitochondria, also contain cholesterol (24). Cholesterol distribution among cellular membranes is governed by a dynamic steady state (25), such that reduction in total cellular cholesterol content by statin or MCD will decrease cholesterol levels in all membranes, including those in the mitochondria (26). Levels of IMM cholesterol can be assessed by the activity of an IMM enzyme, sterol 27-hydroxylase (CYP27A1), which catalyzes the conversion of cholesterol to 27-hydroxycholesterol as a function of cholesterol availability in IMM (27). The amount of 27-hydroxycholesterol in macrophages thus directly reflects IMM cholesterol levels, if CYP27A1 remains constant (26). We found that *Cyp27a1* levels are significantly reduced by 2-day statin treatment (Figure 4-figure supplement 1) but remain steady after 1 h MCD treatment (Figure 4A). We thus analyzed 27-hydroxycholesterol contents by mass spectrometry on MCD-treated macrophages to assess the cholesterol levels in IMM. We find that the amount of cellular 27-hydroxycholesterol was decreased in a dose-dependent manner after a 1 h MCD treatment (Figure 4B), indicating a reduction in cholesterol levels in IMM by MCD. Statins decrease total cholesterol content in macrophages similar to MCD (8), which should then similarly lower cholesterol levels in IMM, even though this particularly assay could not be applied due to the changes in *Cyp27a1* expression described above.

**Figure 4.**
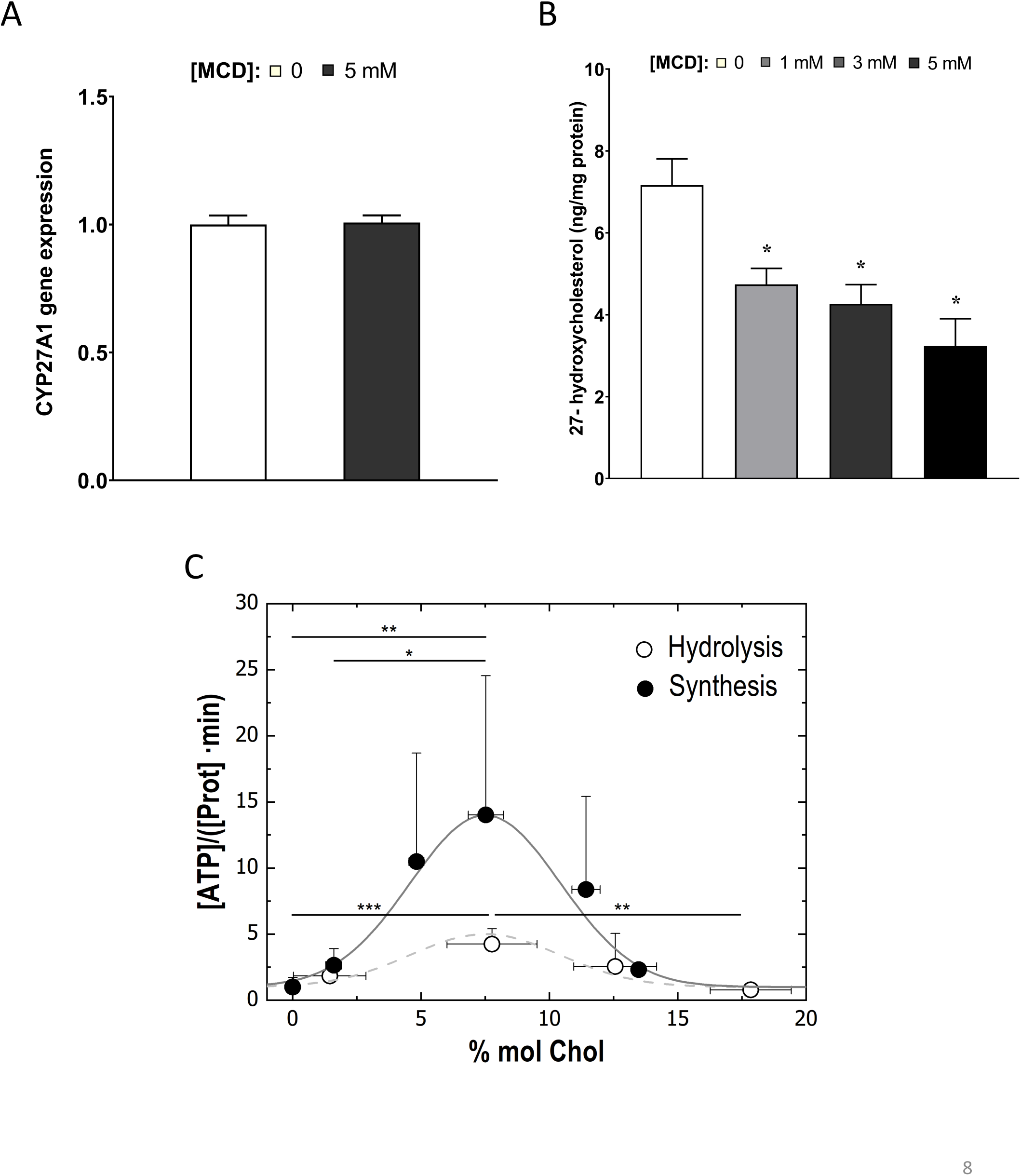
Cholesterol levels modulate the mitochondrial ATP synthase activity. (*A*) sterol 27-hydroxylase, *Cyp27a1*, expression in 1 h MCD-treated and control RAW macrophages. **(*B*)** 27-hydroxycholesterol (27-HC) analysis by ultraperformance liquid chromatography/electrospray ionization/tandem mass spectrometry. 27-HC levels were normalized to the protein content in the whole cell pellet; **(*C*)** ATP hydrolysis and synthesis in cholesterol-doped inner membrane vesicles (IMVs); ATP hydrolysis was performed by adding a total concentration of 2 mM ATP to 200 mM IMVs (lipid concentration) and incubated for 30 minutes. Concentration of phosphates from ATP hydrolysis were measured using the malaquite green assay. ATP concentration after synthesis was measured using ATP detection assay kit (Molecular Probes) with a luminometer GloMax®-Multi Detection. Data in B are from 3 samples per group and data are presented as mean ± SD. An asterisk (*) indicates p<0.05. Data in (C) are representative of at least two independent experiments with three replicates and presented as means± SD. Statistical analysis was performed using the Tukey ANOVA test. (*), (**) and (***) indicate a significant difference with p < 0.05, 0.01 and 0.001, respectively.

To understand precisely how cholesterol levels in the IMM might influence ATP synthase functions, we used an *in-vitro* membrane system composed of mitochondria inner membrane vesicles from *Escherichia coli (*IMVs) (28). *E. coli* inner membrane shares common features with those from mammalian cells (29), including ATP synthase functions (30), but lacks cholesterol or any other sterol derivative in their lipid composition (31). Various levels of cholesterol can be incorporated into the IMVs to assess the impact of cholesterol (32). With this system, we observed that the activity of the ATP synthase, both in synthesis and hydrolysis mode, is highly sensitive to cholesterol concentration in the membrane: the highest activity is found in IMV with membrane containing 7% cholesterol; decreasing cholesterol from 7% cholesterol suppresses ATP synthase activities (Figure 4C). The steady-state IMM cholesterol level in mammalian cells is about 5% (33). If MCD reduces cholesterol in IMM, as we show above (Figure 4B), ATP synthase activity should be suppressed, as we have seen in OCR (Figure 3A, b). Therefore, the *in-vitro* experimental model is consistent with the notion that MCD lowers cholesterol in the IMM, which suppresses ATP synthase activity.

### Reducing cholesterol levels in macrophages alters proton gradients in the mitochondria to activate NF-kB and upregulate Jmjd3

We next studied the potential role of suppressed ATP synthase on NF-κB activation and *Jmjd3* upregulation. ATP synthase in the IMM uses proton flow from the inner space to the matrix to generate ATP (Figure 5A, a). As shown by extracellular flux analysis (Figure 3), MCD suppresses the activity of ATP synthase. This will lead to fewer protons flowing down from the inner space into the matrix and, consequently, more protons will be retained in the inner space in MCD-treated macrophages (Figure 5A, *b*), which could activate NF-κB (34). We tested this using carbonyl cyanide m-chlorophenyl hydrazine (CCCP), a mitochondrial proton ionophore that prevents proton buildup in the inner space (35) (Figure 5A, *c*). Indeed, in the presence of CCCP, MCD failed to activate NF-κB (Figure 5B, *a*). CCCP also prevented MCD from upregulating IL-1β (Figure 5B, *b*). In addition, it abolished *Jmjd3* upregulation by MCD in RAW macrophages and BMDMs (Figure 5C, *a* & *b*), while cells remained fully viable (Figure 5-figure supplement 1A). Moreover, another structurally unrelated mitochondrion proton ionophore BAM15 (36) similarly abolished MCD-induced *Il1b* and *Jmjd3* expression (Figure 5-figure supplement 1B, *a* & *b*). We conclude that reducing cholesterol in macrophages suppresses ATP synthetase activity in the IMM, which likely activates NF-κB and upregulates *Jmjd3*.

**Figure 5.**
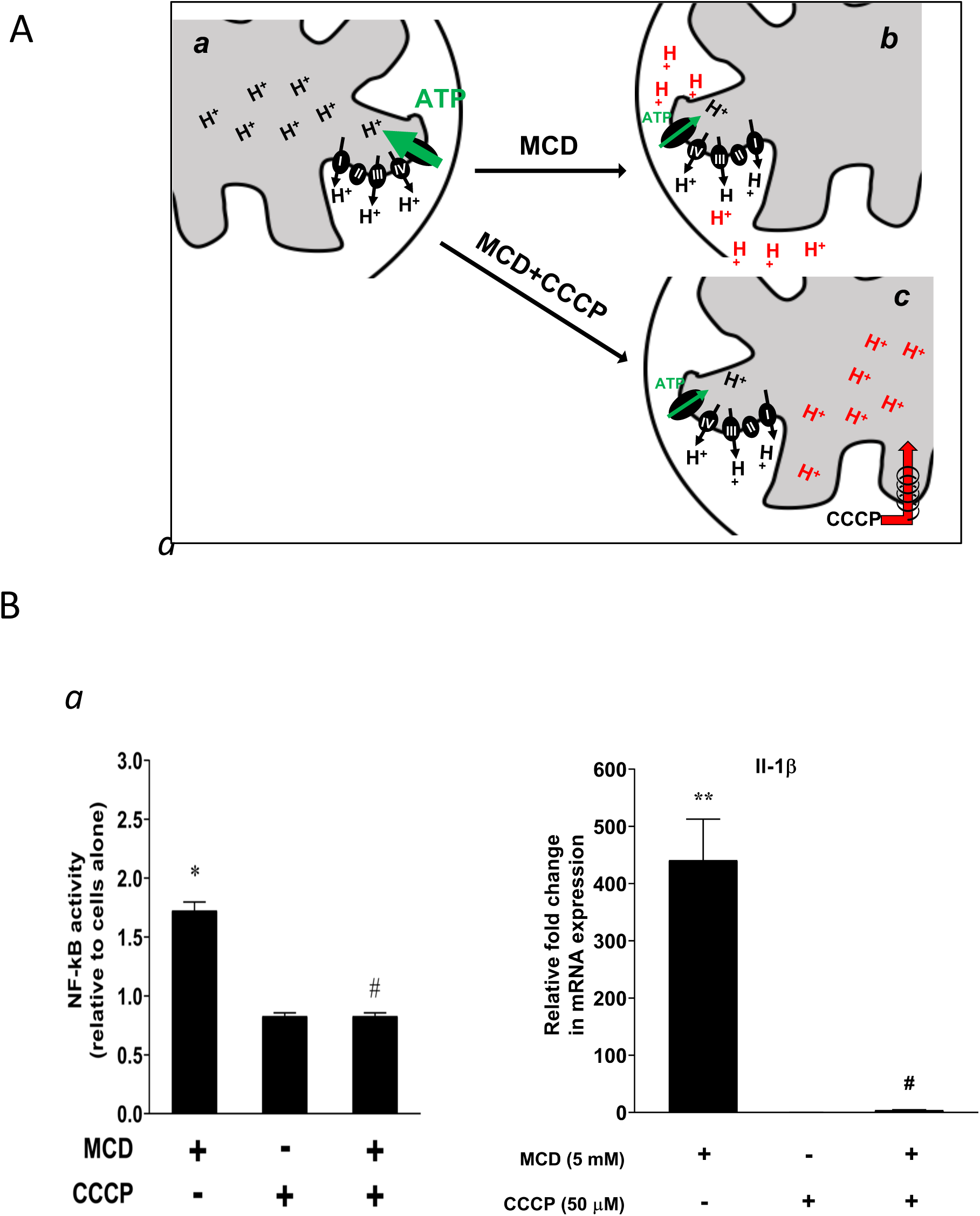
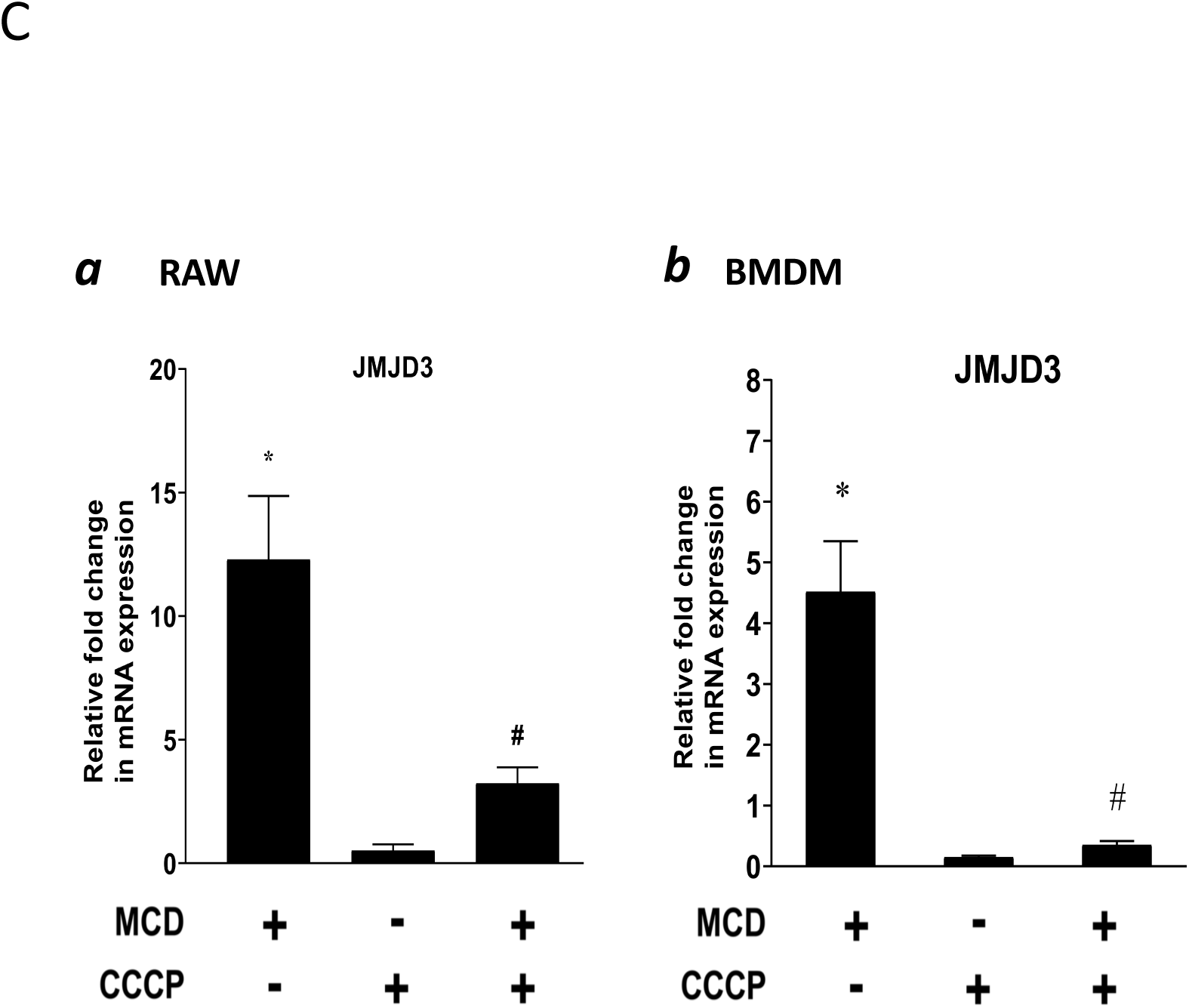
Effect of proton flux on NF-kB activation and *Jmjd3* expression in MCD-treated cells. (***A***) Schematic of potential mechanism induced by MCD, and MCD/CCCP on mitochondrial proton flux. (***B***) Effect of CCCP on NF-kB activity and *Jmjd3* expression: *(a)* on NF-kB activity in RAW blue^TM^ macrophages (CCCP = 50 µM); (*b*) Effect of CCCP on *Il1b* expression in RAW 264.7 macrophages (CCCP = 50 µM). (***C***) Effect of CCCP *(a)* on *Jmjd3* expression in RAW 264.7 macrophages (CCCP = 50 µM) and *(b)* on BMDMs (CCCP = 200 µM). Data are representative of 3 independent experiments with 3 samples per group and data are presented as mean ± SD. Statistical analysis was performed using unpaired, two-tailed Student’s t-test. Asterisk (*) and (**) indicate a significant difference with p<0.05 and p<0.001. A hashtag (#) indicates a significant difference between MCD without or with inhibitors with p < 0.05.

### Reducing cholesterol in macrophages alters chromatin structure

The NF-κB-target gene *Jmjd3* primarily functions to demethylate H3K27me3, an abundant epigenetic mark associated with transcriptional repression (37). The upregulation of *Jmjd3* by statin/MCD is expected to decrease H3K27me3 levels, which should have an impact on the macrophage epigenome. We performed the assay for transposase-accessible chromatin with sequencing (ATAC-seq) to compare the transposase accessibility with or without MCD. We observed that MCD treatment significantly altered the genomic locations of open/close chromatin in macrophages (Figure 6A, a and Supplement file 3). Consistent with our RNA-Seq studies, GSEA of all genes showing altered chromatin accessibility upon MCD treatment identified NF-kB pathway as the top biological process (Figure 6A, b & c). We also analyzed genes being opened by MCD: they predominantly have NF-kB family of TFs in promoters (Figure 6-Figure Supplement 1). We then compared ATAC-seq with RNA-seq and identified overlaps genes. i.e., increased accessibility to transposase and higher expression (Supplement file 4). Noticeably, *Jmjd3*, *Il1b* and *Tnfa* are among those. Also, *Jmjd3* is the only gene with epigenetic modification function. The details of *Il1b* and *Tnfa* is shown in Figure 6B. We conclude that the epigenome is altered by cholesterol reduction in macrophages.

**Figure 6.**
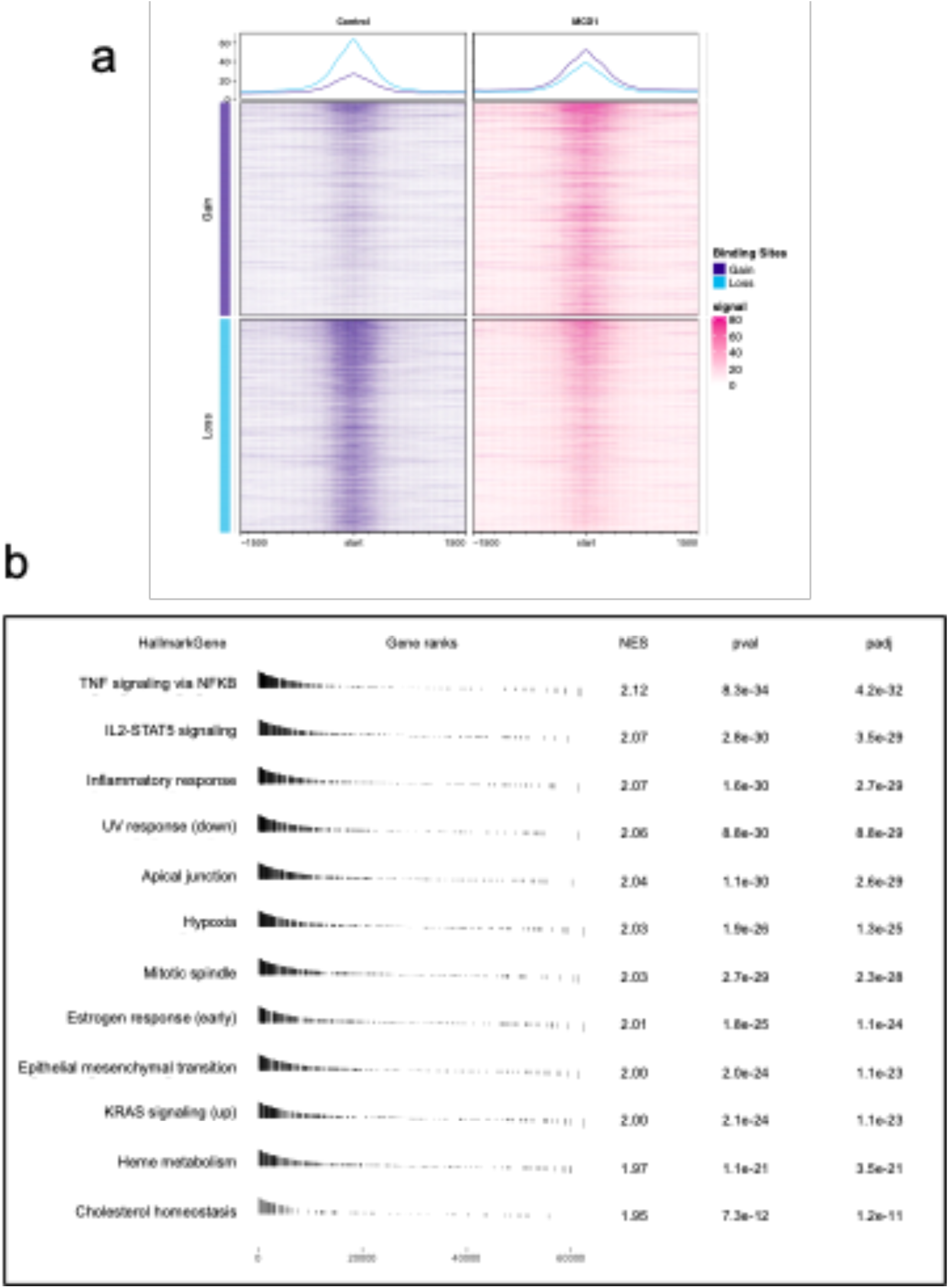
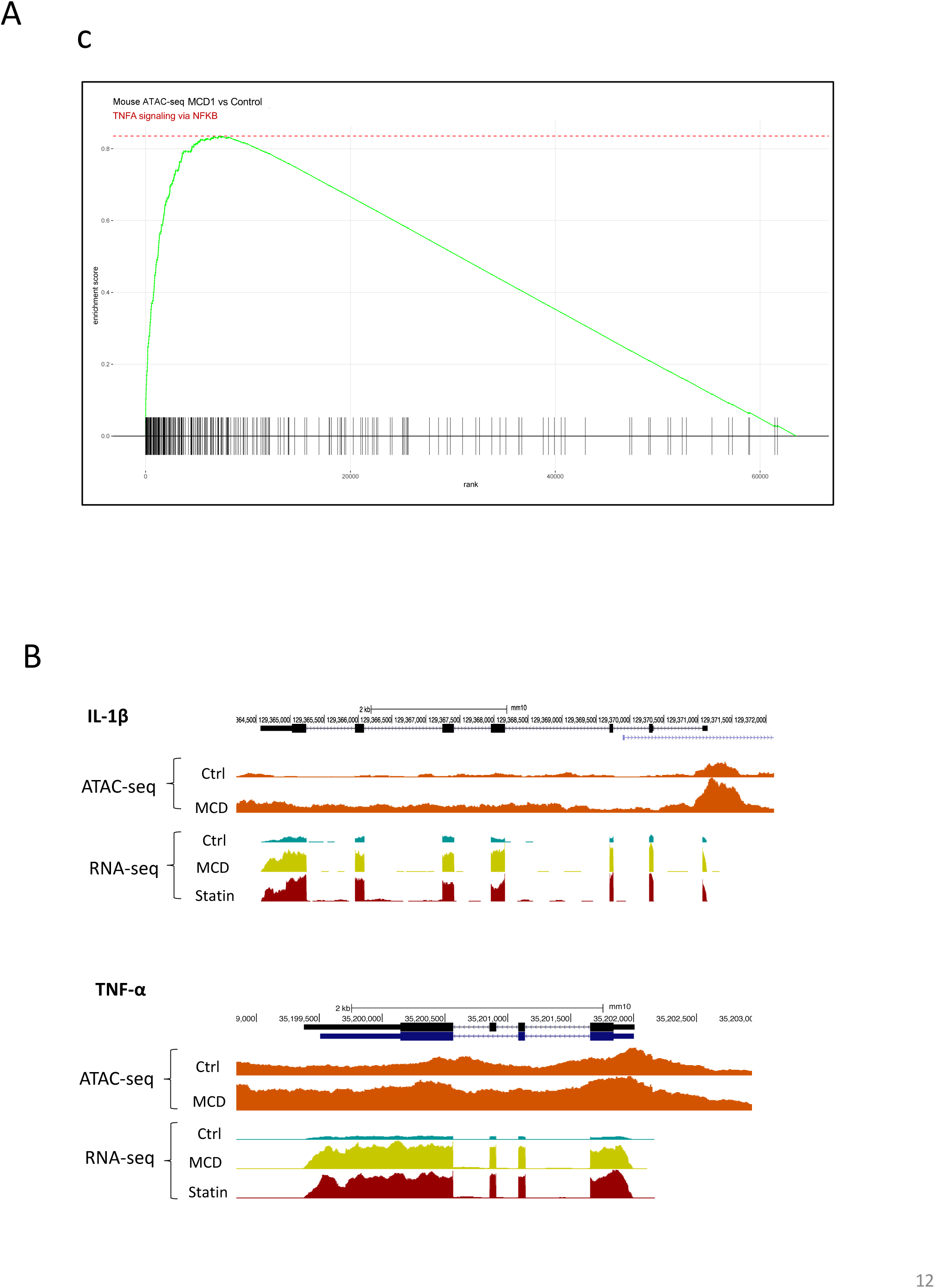
Cholesterol modulates macrophage epigenetic modifications. (*A*) ATAC-seq in control and MCD (5 mM, 1 h) treated RAW 264.7 macrophages. *(a)* Summit-centered heatmap of differentially accessible ATAC-seq signals. *(b)* Pathways identified by GSEA of differentially assessable genes in *(a)*. *(c)* The details of most highly represented pathway, TNFA signaling vis NFKB. **(*B*)** ATAC-seq and RNA-seq profiles alignment from ATAC-seq and RNA-seq for the genomic loci of *Il1b* (top) and *Tnfa* (bottom).

### Reducing cholesterol promotes anti-inflammatory responses in activated macrophages

The reconfigured epigenome upon cholesterol depletion could poise resting macrophages for different inflammatory responses. We therefore tested inflammatory responses in statin-treated macrophages with LPS (classically activated or M1 phenotype) or with IL-4 (alternatively activated or M2 phenotype). Macrophages were treated with statins or without and then stimulated by LPS. RNA-seq was performed. Cholesterol reduction by statins altered a large number of genes, up or down, in LPS-stimulated macrophages (Figure 7A, *a*; Supplementary file 5). Gene Ontology (GO) analysis of genes with decreased expression upon statin treatment revealed that statins primarily suppress inflammatory processes (Figure 7A, *b*), while genes involved in cellular homeostatic functions were upregulated (Figure 7A, *c*). More specifically, in BMDMs activated by LPS, statin treatment led to suppressed expression of pro-inflammatory cytokines (*Il1b*, *Tnfa*, *Il6* and *Il12*), but enhanced expression of anti-inflammatory cytokine *Il10* (Figure 7B). When BMDMs were activated alternatively by IL-4, statin treatment promoted the expression of IL-4 target genes *Arg1*, *Ym1* and *Mrc1* (Figure 7C). The enhanced expression of the anti-inflammatory genes (*Arg1*, *Ym1*, *Mrc1* and *Il10*) by statins in activated macrophages, both M1 and M2, was also correlated with the removal of H3K27me3, a repressive marker and the substrate of JMJD3 in resting macrophages (Figure 7D). Thus, experimental evidence supports the notion that cholesterol reduction by statins in macrophages leads to less pro-inflammatory responses to LPS, but higher expression of anti-inflammatory genes, such as these activated by IL4, correlated with H3K27me3 removal (38).

**Figure 7.**
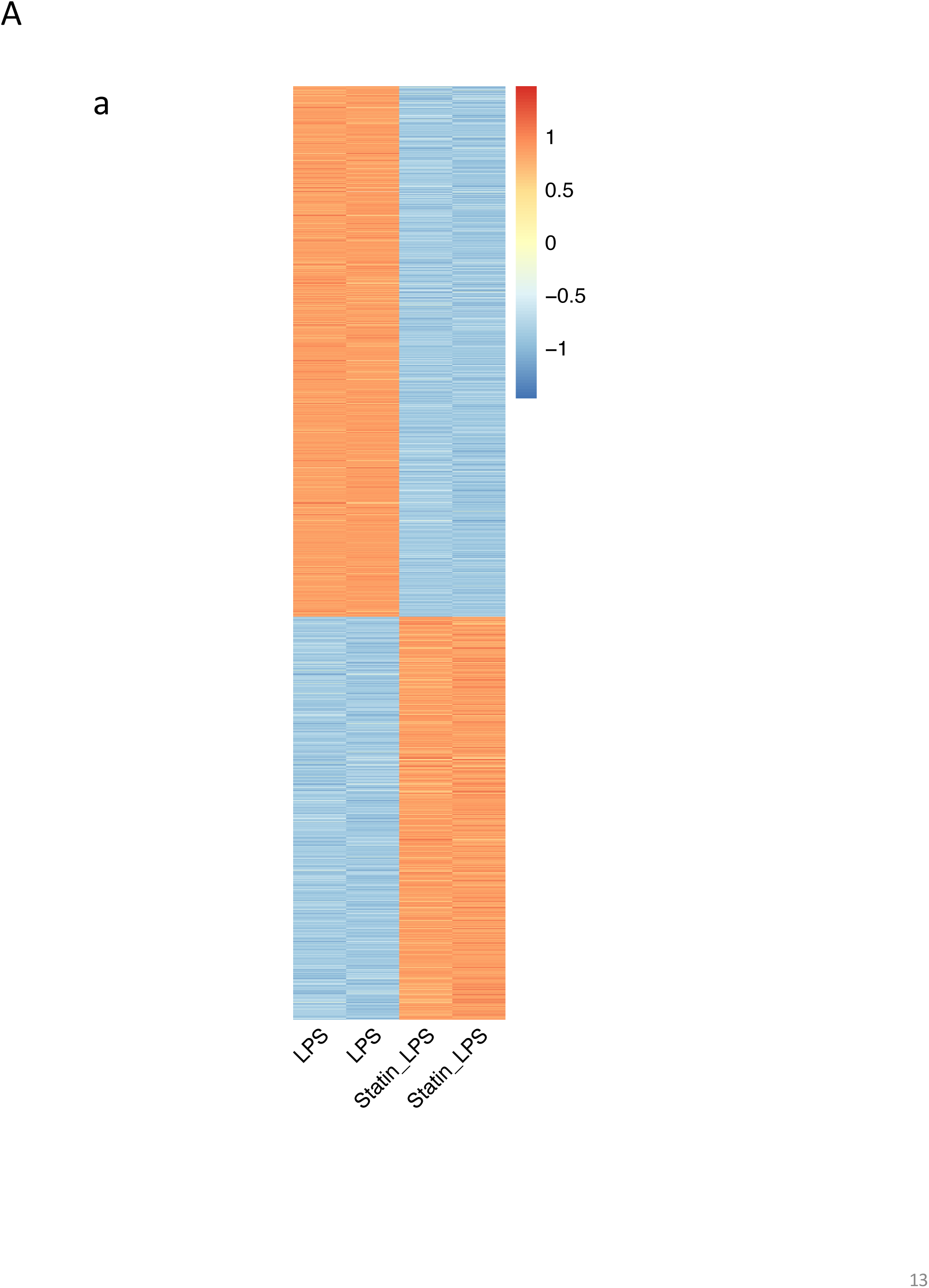
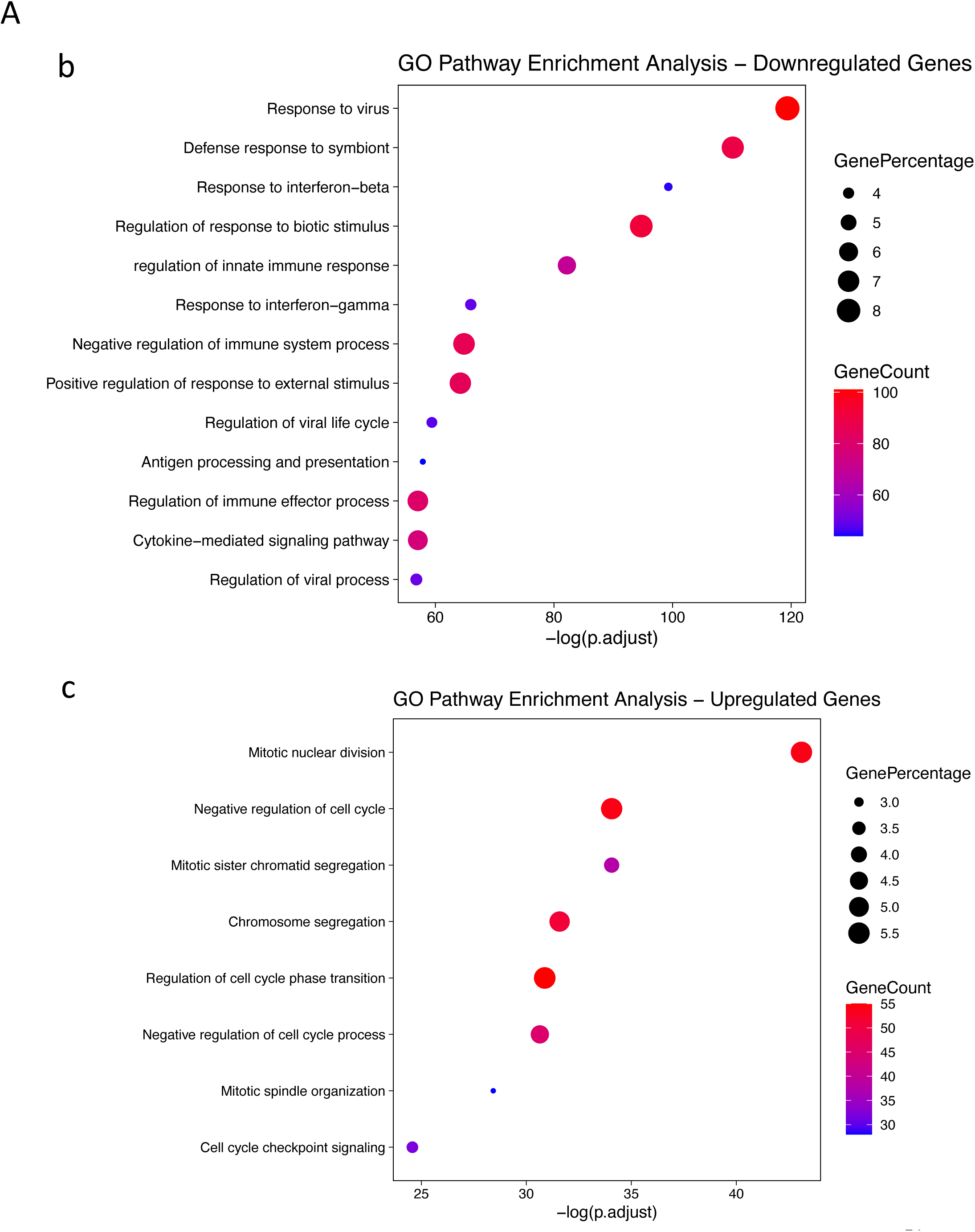
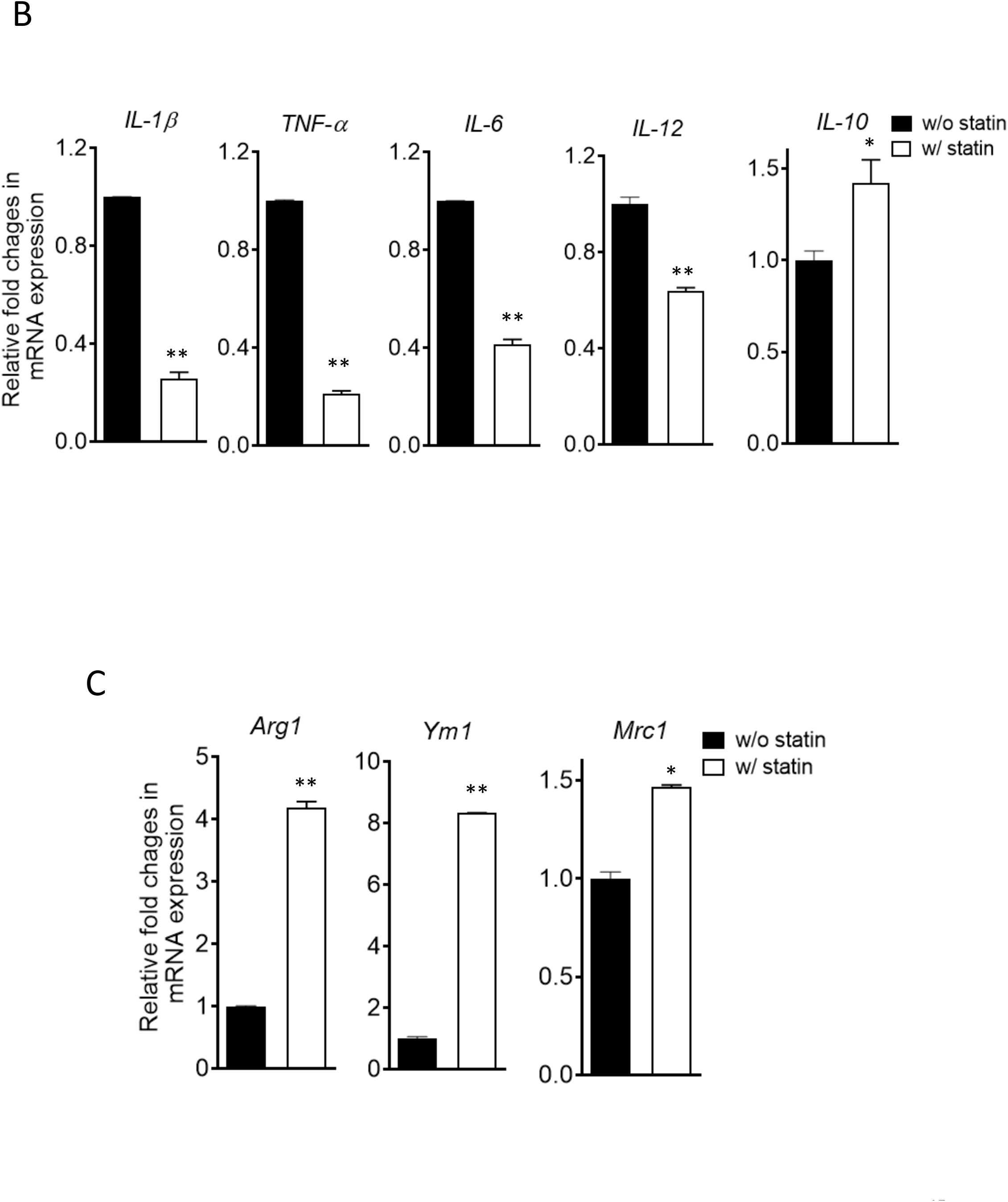
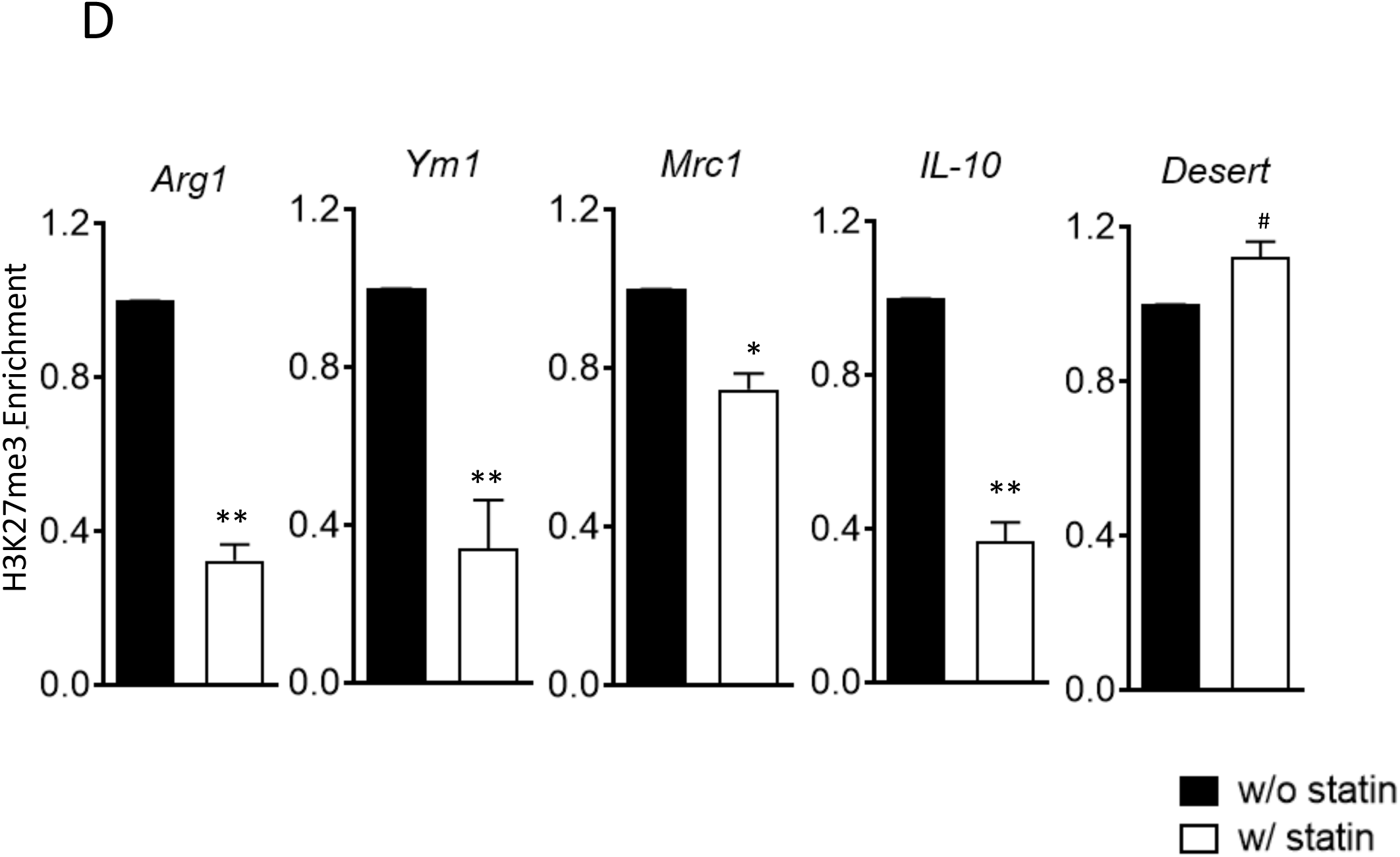
Statins supress proinflammatory cytokines and enhance anti-inflammatory factors in LPS or IL-4 activated macrophages. *(A)* RAW 264.7 macrophages treated with statin (lovastatin, 7 µM + 200 µM mevalonate; 2 days) or without were stimulated with LPS (100 ng/ml) for 3 h. *(a)* Heatmap of differentially expressed genes by statin; *(b)* GO analysis of down-regulated genes by statin. *(c)* GO analysis of up-regulated genes by statin. **(*B*)** BMDMs treated with or without compactin (10 µM + 200 µM mevalonate; 2 days) are stimulated by LPS (50 ng/ml, 3 h). Gene expressions are analyzed by qPCR. ***(C)*** BMDMs treated with or without compactin are stimulated by IL-4 (20 ng/ml, 6 h) and gene expressions are analyzed by qPCR. (***D***) Chromatin-immunoprecipitation (ChIP) analysis of H3K27me3 in BMDMs with or without compactin treatment. The inactive gene *desert* is used as input control. Data are representative of at least 2 independent experiments with 3 samples per group and data are presented as mean ± SD. Statistical analysis was performed using unpaired, two-tailed Student’s t-test. An asterisk (*) and (**) indicate a significant difference with p<0.05 and p<0.001. A hashtag (#) indicates not significant.

### Anti-inflammatory responses by cholesterol reduction in macrophages rely on Jmjd3 and its demethylase activities

*Jmjd3* belongs to the JmjC demethylase family that requires α-ketoglutarate (α-KG) as co-factor to demethylate histone (38). Noticeably, *Jmjd3* (*Kdm6b*) is the only gene member of the JmjC family upregulated by MCD in macrophages and, importantly, the expression of closely related *Utx* (*Kdm6a*) is not changed by cholesterol reduction (Figure 8A, a). This presented an opportunity to specifically probe the involvement of *Jmjd3* demethylation activity in suppressing *Tnfa* yet raising *Il10* expression by MCD. *Jmjd3* expression is increased with MCD in a concentration-dependent manner (Figure 8-figure supplement 1). When subsequently stimulated by LPS, MCD-treated macrophages dose-dependently express less *Tnfa*, but more *Il10*: the ratio of *Il10*/*Tnfa* rises with MCD concentrations (Figure 8A, *b*, white bars). However, if glutamine, the precursor of α-KG (38), is absent in the medium, there is little change in the ratio of *Il10*/*Tnfa* (Figure 8A, *b*, black bars), regardless of MCD concentration. Furthermore, if the demethylase activity of Jmjd3 is inhibited by a specific inhibitor GSKj4 (39), the ratio of *Il10*/*Tnfa* also failed to rise upon MCD treatment (Figure 8A, *c*). BMDMs similarly modify their response upon glutamine availability. When stimulated by LPS, MCD-treated BMDMs rise *Il10*/*Tnfa* ratio, dependent on glutamine: *Il10*/*Tnfa* fails to increase by MCD in the absence of glutamine (Figure 8B, *a*). Glutamine is also necessary for MCD to boost IL4-targeted *Arg1* (Figure 8B, *b*). We next used shRNA to knockdown *Jmjd3* (*Jmjd3* KD) (Figure 8-figure supplement 2) and tested its impact on statin-treated macrophages. When activated by LPS, statin-treated *Jmjd3* KD macrophage failed to raise *Il10*/*Tnfa* ratio (Figure 8C, a). *Jmkd3* KD also abolished the rise of *Arg1* by statin when stimulated by IL-4 (Figure 8C, b). Together, we conclude that *Jmjd3* and its demethylase activity are necessary to promote the expression of anti-inflammatory elements upon cholesterol in macrophages.

**Figure 8:**
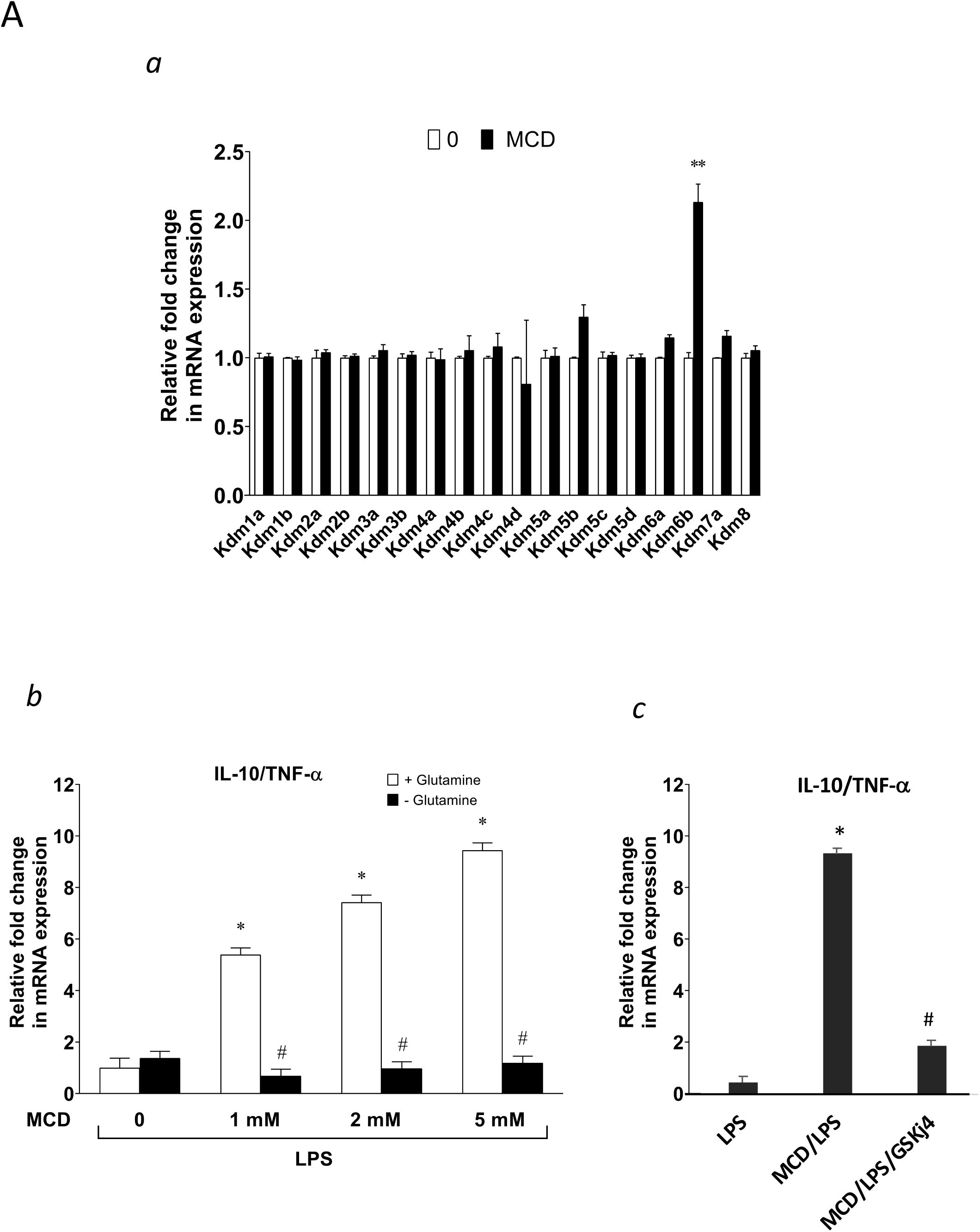
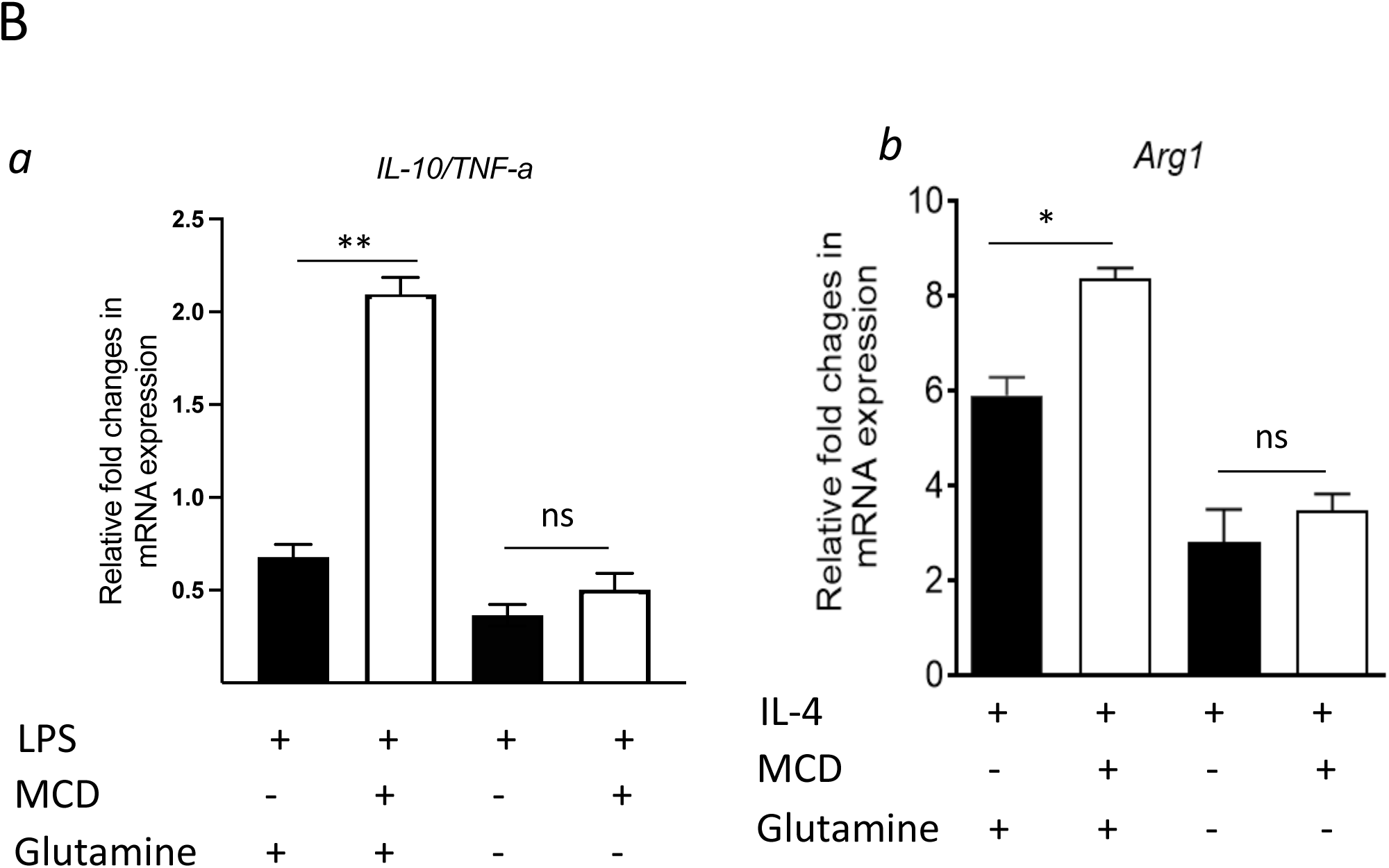
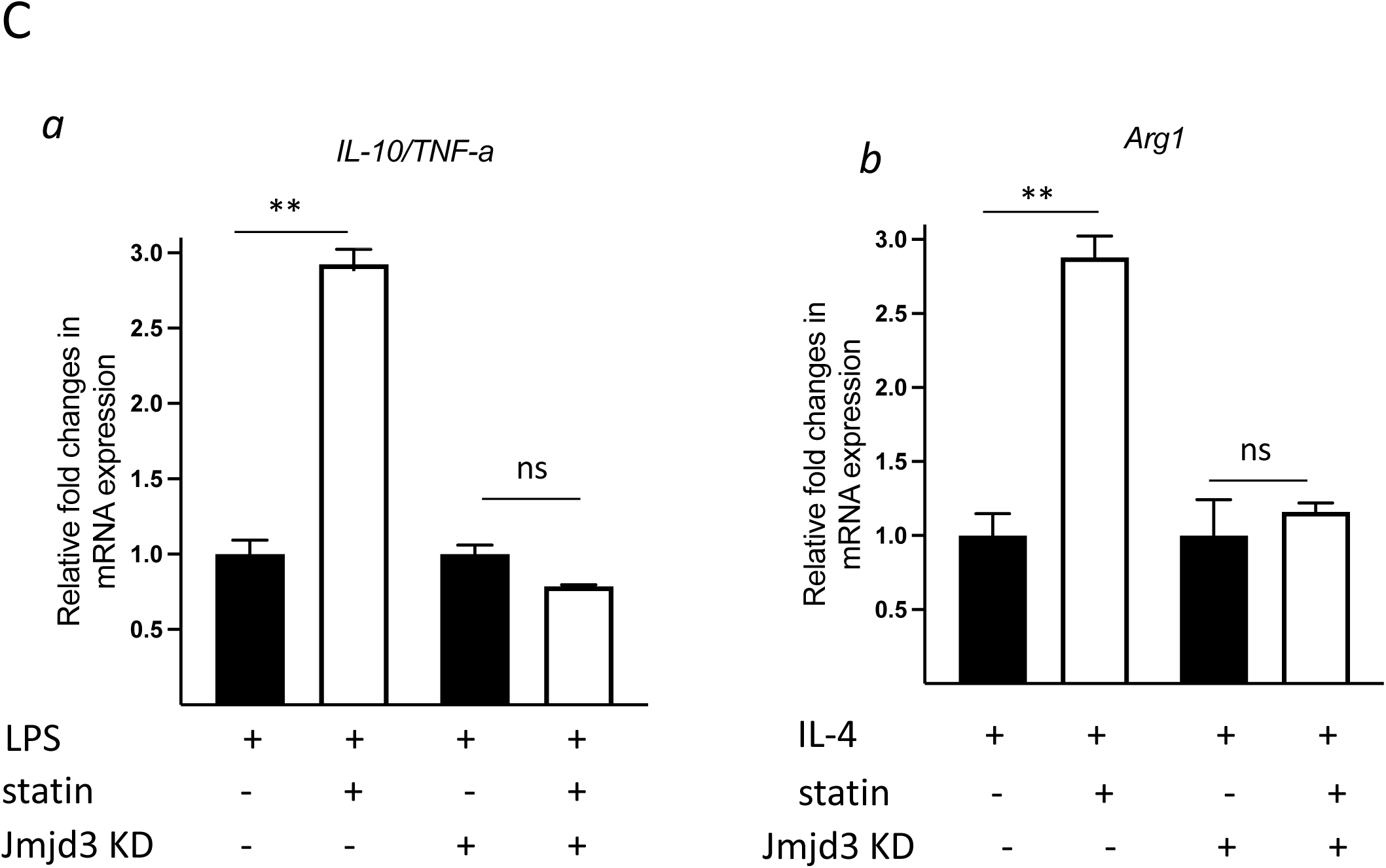
Cholesterol reduction suppresses proinflammatory phenotypes and enhance the expression of anti-inflammatory factors, depending on *Jmjd3* and its enzymatic activity. *(A)* RAW macrophages were treated with MCD (5 mM, 1 h). *(a)* The expression of JmjC demethylase family with or without MCD treatment. *(b)* After LPS stimulation (100 ng/ml, 3 h), in the presence of glutamine or without, the expression of *Il10* and *Tnfa* were analyzed by qPCR to generate the ratio of *Il10*/*Tnfa*. *(c)* Effect of GSJK4, a JMJD3 inhibitor, on *Il10*/*Tnfa*. (*B*) *(a)* BMDMs treated with or without MCD are stimulated by LPS (50 ng/ml, 3 h) in the presence of glutamine or without. Expression of *Il10* and *Tnfa* were analyzed by qPCR to generate the ratio of ratio of *Il10*/*Tnfa*. *(b)* BMDMs treated with or without MCD are stimulated by IL-4 (20 ng/ml, 6 h) and expression of *Arg1* is analyzed by qPCR. *(C)* wt and *Jmjd3* KD RAW macrophages were treated with compactin (10 µM + 200 µM mevalonate; 2 days) and then stimulated with LPS (100 ng/ml, 3 h) or IL-4 (20 ng/ml, 6 h). Expression of *Il10* and *Tnfa (a)* or *Arg1 (b)* were analyzed by qPCR. Data are representative of 3 independent experiments with 3 samples per group and data are presented as mean ± SD. Statistical analysis was performed using unpaired, two-tailed Student’s t-test. An asterisk (*) and (**) indicate a significant difference with p<0.05 and p<0.001. A hashtag (#) indicates a significant difference between MCD without or with glutamine with p < 0.05.

### Statin treatment in vivo also reduces cholesterol content, upregulates Jmjd3, and promotes anti-inflammatory gene expression in macrophages

To test the *in vivo* effect of statins on macrophages, mice were fed with statins or not for 14 days and the peritoneal macrophages were isolated (40) and tested for inflammatory responses. The cholesterol content was decreased by about 20% in freshly isolated peritoneal macrophages from statin-fed mice, compared to those from control animals (Figure 9A). Expression of *Jmjd3* was upregulated by statin-feeding (Figure 9B). When subsequently challenged by LPS, macrophages from statin-fed mice showed lower expression of pro-inflammatory cytokines (*Il1b*, *Tnfa*, *Il6* and *Il12*), and enhanced expression of anti-inflammatory cytokine *Il10*, relative to those from controls (Figure 9C). In addition, when activated alternatively by IL-4, macrophages from statin-fed mice expressed higher levels of Arg*1*, *Ym1* and *Mrc1*, compared to these from controls (Figure 9D). Thus, statin treatment *in vivo* decreases cholesterol content, upregulates *Jmjd3,* and promotes anti-inflammatory functions in freshly isolated peritoneal macrophages, in an identical fashion to that of BMDMs treated *in vitro* with statins (Figure 7B&C).

**Figure 9:**
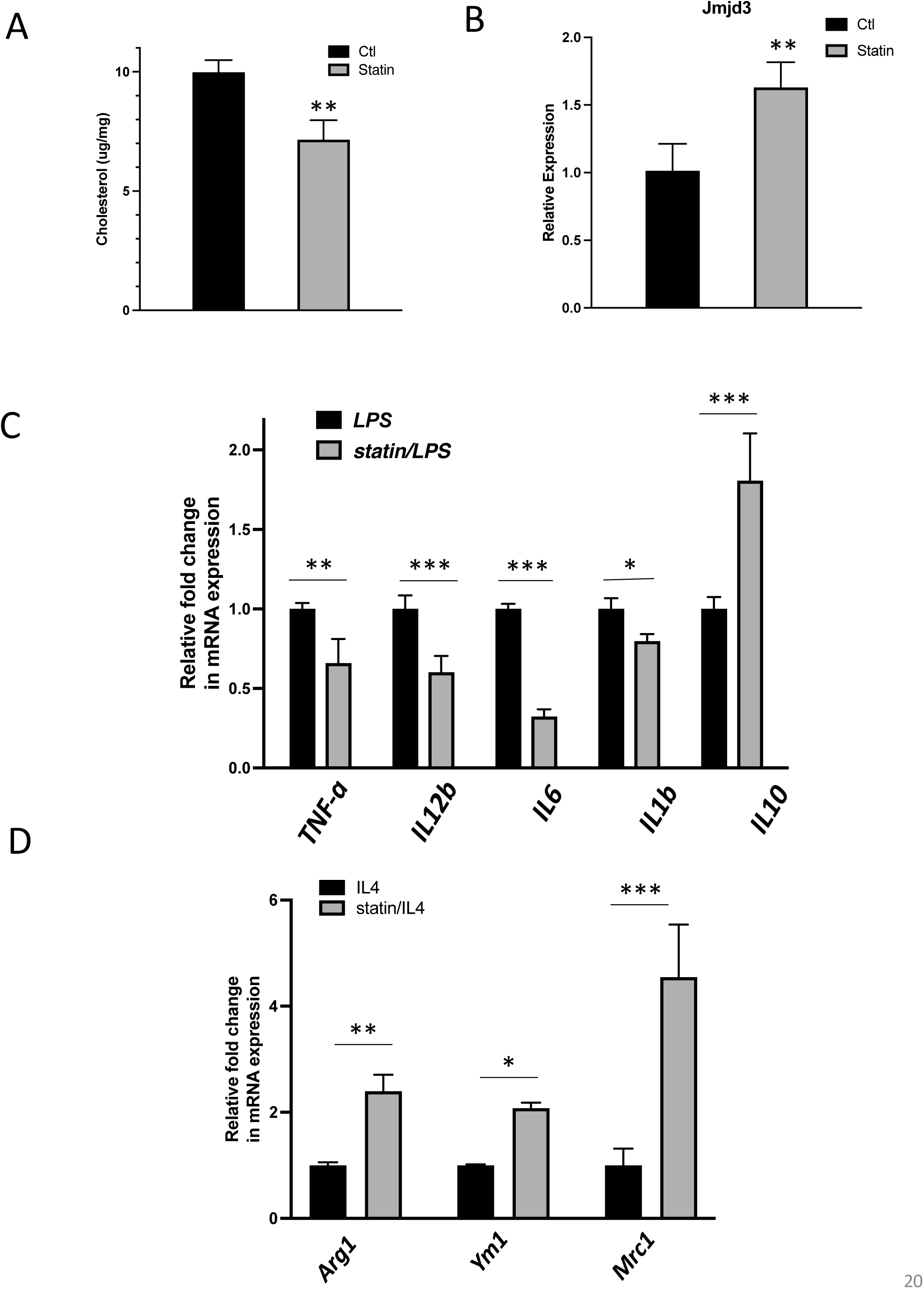
*In vivo* statin-feeding in mice reduces cholesterol and upregulates *Jmjd3* in the peritoneal macrophages, which conveys anti-inflammatory phenotype. *(A)* Total cholesterol contents (free and esterified) in freshly isolated mouse peritoneal macrophages from simvastatin-fed (100 µg/kg/day, 14 d) and control mice. (*B*) *Jmjd3* gene expression in peritoneal macrophages from simvastatin-fed (statin) and control mice. (*C*) Freshly isolated mouse peritoneal macrophages from simvastatin-fed and control mice are stimulated by LPS (100 ng/ml, 6 h). Gene expressions are analyzed by qPCR. **(D)** Freshly isolated mouse peritoneal macrophages from simvastatin-fed and control mice are stimulated by IL-4 (20 ng/ml, 6 h) and gene expressions are analyzed by qPCR. Data are representative of 3 independent experiments with 3-4 samples per group and presented as mean ± SD. Statistical analysis was performed using unpaired, two-tailed Student’s t-test. An asterisk (*), (**) and (**) indicate a significant difference with p<0.05, p<0.005 and p<0.001, respectively.

## DISCUSSION

In this study, we show that reducing cholesterol in macrophages, either with statins or MCD, upregulates Jmjd3 through suppressing mitochondria respiration. Cholesterol reduction also modifies the epigenome in macrophages. Upon subsequent activation by either M1 or M2 stimuli, statin-treated macrophages are phenotypically more anti-inflammatory. This anti-inflammatory phenotype is also observed in peritoneal macrophages freshly isolated from statin-treated mice.

We speculate that *Jmid3* is the key responsible factor. Macrophages have two H3K27me3 demethylases, *Utx* and *Jmjd3*. Only *Jmjd3* is upregulated by statin or MCD. It is plausible that *Jmjd3*, by removing H3K27me3 from *ll10*, *Arg1*, *Yam1*, and *Mrc1*, poises these genes in resting macrophages to promote their activated expression. On the other hand, statins suppress a large group of proinflammatory genes activated by LPS, which could not directly be attributed to *Jmjd3*. However, *Il10* is poised by Jmjd3 and shows higher expression upon LPS stimulation. It could be that upregulation of *Il10* leads to the suppression of the proinflammatory phenotype under LPS activation. Recent studies have shown that endogenously produced IL-10, through autocrine/paracrine mechanisms, modulates mitochondria respiration to inhibit cellular events, such as glycolysis and mTORC1. This suppresses the expression of proinflammatory genes (41, 42). The elevated *Il10* expression in statin/MCD-treated macrophages could play a similar role to suppress the proinflammatory phenotypes. Of note, this current study focuses mostly on the regulation of gene expression. The ultimate impact of statins on inflammation should be confirmed at the protein level in future studies.

We also document that reducing the level of macrophage cholesterol alters the epigenome, concurrent with *Jmjd3* upregulation. Removing H3K27me3 by JMJD3 could open certain genome regions and contribute to the changes in the epigenome seen in ATAC-seq. However, the changes we observed are much more profound, indicating that other epigenetic modifiers, are activated by cholesterol reduction. Future studies will be required to identify these modifiers. Nevertheless, several lines of evidence support the notion that *Jmjd3* is most relevant to inflammatory activation. First of all, only *Jmjd3*, among the members of the JmjC demethylase family, is upregulated by statin or MCD. *Utx*, also a H3K27me3 demethylase, is not altered by statin or MCD. Secondly, the JmjC demethylase family members require glutamine, the source of a-ketoglutarate, for demethylation. Since *Jmjd3* is the only upregulated demethylase, glutamine could be used to specifically probe *Jmjd3* demethylation function in macrophages M1/M2 activation. Glutamine is indeed necessary for MCD to modulate both LPS and IL-4 activation. Thirdly, GSKj4, the inhibitor for H3K27me3 demethylases (Utx and Jmjd3), abolishes the MCD effect. Furthermore, knockdown of *Jmjd3* by shRNA diminishes the effect of statins on LPS or IL-4 activations. Thus, the upregulation of *Jmjd3* by statin/MCD significantly contribute to poise the epigenome in resting macrophages, which controls the subsequent inflammatory response to LPS or IL-4.

Our results here support the notion that the epigenome in resting macrophages is largely poised for inflammatory activation (43). This also agrees with previous studies where cholesterol reduction was shown to decrease proinflammatory responses to multiple stimuli against multiple TLRs (9). We have focused on classically activated (M1) and alternatively activated (M2) phenotypes, two extremes of inflammatory activation in macrophages. This could be an experimental starting point, since macrophages likely encounter a wide range of stimuli within a continuum between M1 and M2 phenotypes. The poised epigenome in resting macrophages, i.e., prior to activation, may serve as an initial blueprint to propel the activated inflammatory responses. In addition, the inflammation processes are thought to initially engage the M1 phenotype to fight pathogens and then gradually gain the M2 phenotype to restrain excessive damage and restore tissue homeostasis (15).

Another novel finding here is that the level of cellular cholesterol directly controls mitochondria respiration. Except for specialized cell types (i.e., steroidogenic cells), the mitochondrial cholesterol in mammalian cells is in equilibrium with overall cellular cholesterol and, as such, fluctuates with cellular cholesterol through a dynamic steady state (24). Our study here for the first time suggests the mitochondrial membrane as a locus sensing the cellular cholesterol level, which in turn contributes to the regulation of metabolic processes and gene expression. This is somewhat analogous to the regulation of Sterol regulatory element-binding proteins (SREBP) pathways by the cellular cholesterol level through the endoplasmic reticulum (ER) membrane (44). We speculate that the mitochondria likely function far beyond the traditionally called powerhouse that produces ATP (45).

Macrophages are a major component in both the innate and adaptive immune systems. The anti-inflammatory effect of statins is essentially due to their primary pharmacological action discovered in 1970s, I.e., the inhibition of HMG-CoA reductase (46). This decreases the level of circulating LDL and as well as inhibits *de novo* synthesis of cholesterol in the macrophages themselves. Macrophages with less cholesterol are anti-inflammatory, thereby contributing to the anti-inflammatory action of statins.

## Data availability

Sequencing data have been deposited in GEO under accession codes: GSE196187, GSE196188, GSE196189. All data generated or analysed during this study are included in the manuscript.

## Acknowledgements

This work was supported by a grant (grant-in-aid) from Heart and Stroke Foundation of Canada (HSFC), G-19-0026359, and a grant from Canadian Institute of Health Research (CIHR), PJT-180504. This work was also supported in part by the TECNOLOGÍAS 2018 program funded by the Regional Government of Madrid (Grant S2018/BAA-4403 SINOXPHOS-CM, to ILM) and by a grant from NIH (HL020948, to JGM). We thank Dr. T. Lagace for reagents, and Dr. M. Harper for access to use the XFe96 Seahorse Extracellular Flux Analyzer. We also thank the Ottawa Hospital Research Institute’s Biotherapeutic Manufacturing Centre-Virus Manufacturing Facility for making shRNA lentivirus. We knowledge the technical assistance of H. Bandukwala, Yuefeng Li and E. S. Qamsar.

## Materials and Methods

### Reagents and chemicals

Cell culture Dulbecco’s Modified Eagle Medium (DMEM) was purchased from Gibco (Thermo Fisher Scientific, Watham, MA). Antibiotics (penicillin and streptomycin) and fatty acid (FA)-free bovine serum albumin (BSA) were purchased from Sigma-Aldrich (St Louis, MO). FBS (Optima) was purchased from Atlanta Biologicals (R&D systems; Minneapolis, MN). As for the chemicals, the following inhibitors: MG-132 (proteasome inhibitor), BAY11-7082 (IKK inhibitor), and BAM15 (another mitochondrial uncoupler) were purchased from TOCRIS chemicals (part of R&D systems). Methyl-β-cyclodextrin (MCD), simvastatin, mevalonate, oligomycin, rotenone, antimycin A, carbonyl cyanide 3-chlorophenylhydrazone (CCCP), 4-(2-hydroxyethyl)-1-piperazineethanesulfonic acid (HEPES), and phosphate-buffered saline (PBS) were purchased from Sigma-Aldrich. For the IMVs study, magnesium chloride (MgCl_2_), 2-(N-morpholino ) ethanesulfonic (MES), sucrose, hexane, bovine serum albumin (BSA), sulfuric acid (H_2_SO_4_), hydrochloric acid 37% (HCl), sodium chloride (NaCl), potassium chloride (KCl), dibasic potassium phosphate (K_2_HPO_4_), sodium hydroxide (NaOH), Hydrogen peroxide (H_2_O_2_), trichloroacetic acid (TCA), adenosine 5ʹ-triphosphate disodium salt hydrate (Na_2_ATP), adenosine 5ʹ-diphosphate sodium salt (Na_2_ADP), and cholesterol were also supplied by Sigma-Aldrich. Ammonium molybdate (VI) tetrahydrate and L-ascorbic acid were purchased from Acros Organics (part of Sigma Aldrich). Ultrapure water was produced from a Milli-Q unit (Millipore, conductivity lower than 18 MΩ cm). Rabbit anti-mouse JMJD3 primary antibody was lab-generated as described in^1^. The following antibodies were acquired from several vendors: mouse monoclonal anti-β-actin antibody (A1978) from Sigma-Aldrich, rabbit anti-mouse pCREB (87G3), rabbit anti-mouse CREB (86B10), and rabbit anti-mouse Tri-Methyl-Histone H3 (Lys27) antibodies were from Cell Signaling technology (Danvers, MA). The secondary antibody HRP-conjugated anti-rabbit antibody was from Cayman chemicals (Ann Arbor, MI). The Enhanced chemiluminescence (ECL) solutions for the Western blotting system were from GE Healthcare (Chicago, IL). The protease and phosphatase inhibitor cocktails were purchased from Sigma-Aldrich.

### RAW264.7

RAW 264.7 TIB-71 cells were directly ordered from American Type Collection Culture (ATCC). Cells were tested for “Mycoplasma contamination” by ATCC Cell Authentication Service in October 2023 and result was negative. Cells was maintained in 100-mm diameter tissue culture-treated polystyrene dishes (Fisher Scientific, Hampton, NH) at 37°C in a humidified atmosphere of 95% air and 5% CO_2_. The cells were cultured in DMEM-based growth medium containing glutamine and without pyruvate, supplemented with 10% [v/v] heat-inactivated fetal bovine serum (FBS) and 1x Penicillin/ streptomycin). For experiments (and routine subculture), cells were collected in growth media after 25 minutes incubation with Accutase (Sigma Aldrich, St Louis, MO) at 37°C in a humidified atmosphere of 95% air and 5% CO_2_. In preparation for experiments, cells were seeded in 6-well tissue culture-treated polystyrene plates (Fisher Scientific, Hampton, NH) at 30,000 cells/cm^2^ for 2 days until cells have recovered in time for treatments. Unless otherwise indicated, cell treatments were prepared in DMEM supplemented with 0.25% (w/v) FA-free BSA that was sterilized by filtration through 0.2-µm pore size cellulose acetate syringe filters. For treatments, cells were washed with pre-warmed sterile PBS and were treated with media containing MCD (5 mM), cell culture-grade water (negative control). Then, the cells were incubated for 1 hour under cell culture conditions. RAW 264.7 macrophages were pre-incubated for 5 minutes with CCCP (50 µM), 30 minutes with BAM15 (200 µM), or 1 hour with NF-kB inhibitors (MG-132 [5 µM] or BAY11-7082 [10 µM]).

### Bone marrow-derived macrophages (BMDMs)

Bone marrow-derived macrophages (BMDM) were differentiated from bone marrow cells isolated from the femora, and tibiae of 4- to 16-week-old wild type C57BL/6J mice. Euthanasia was performed by CO_2_ gas asphyxiation followed by cervical dislocation. Euthanized mice were soaked with 70% (v/v) ethanol immediately prior to dissection. After careful isolation of the bones, the ends of the bones were cut with sterile scissors and centrifuged at 10,000 rpm for 15 seconds at room temperature in a microcentrifuge (MCT) tube where the bone marrow is collected. The bone marrow is then filtered using sterile, combined and centrifuged at 480 *g* for 10 minutes at room temperature. The cell pellet is resuspended briefly (< 2 minutes) in red blood cell lysis buffer. The differentiation medium (DMEM + 20% L-929-conditioned media + 10% FBS + 1% penicillin-streptomycin) is then added and cells are centrifuged again at 480 *g* for 10 minutes at room temperature. The cell suspension is filtered once more through a 70-µm strainer (into another 50 ml centrifuge tube to rinse and collect all the cells. 100-mm diameter suspension dishes (Greiner Bio-One, Monroe, NC) are seeded with 10 mL per dish at 0.6-0.8 x 10^6^ cells/mL (∼10,000-15,000 cells/cm^2^). The dishes were then incubated for 6 days at 37°C in a humidified atmosphere of 95% air and 5% CO_2_. Differentiation media (10 mL) are added on Day 3 or 4.

At the end of day 7, cells are detached with trypsin, counted and reseeded into 6-well plates at 0.5 million cells/well (∼ 50,000 cells/cm^2^) using 2mL/well of DMEM supplemented with 10% FBS and 1% P/S. Plates are left overnight at 37°C in a humidified atmosphere of 95% air and 5% CO_2_ to ensure cell adherence. BMDMs are now ready to be used for assays.

### JMJD3 shRNA preparation

For the JMJD3-shRNA construct, the ShKDM6B-265-Up CCGG CCTCTGTTCTTGAGGGACAAA CTCGAG TTTGTCCCTCAAGAACAGAGG TTTTTG and ShKDM6B-265-Down AATTCAAAAA CCTCTGTTCTTGAGGGACAAA CTCGAG TTTGTCCCTCAAGAACAGAGG sequences were used to generate lentivirus harboring shRNA or empty Neo (control). The shRNAs were cloned in lab into pLKO.2 Neo plasmid using EcoR1 and AgeI restriction enzymes. The JMJD3 shRNA lentiviral titer was prepared using a polyethylenimine (PEI)-based transfection. Briefly, on Day 1 HEK 293T cells were seeded at 8 x 10^6^ cells in 15 cm dishes. On Day 2 (transfection day), before the start of transfection, media was removed, cells were washed with media/PBS, then fresh media (media containing 5% FBS) is added. 30µg total DNA was used (17.5 µg of shRNA) to transfect the cells seeded. OptiMEM media was used to prepare the transfection mix where 1 ml DNA is mixed to 1 ml PEI reagent, and incubated for another 20-30 min. The mix is added gently on to the cells, drop by drop. After overnight incubation, the media is replaced and fresh 5 % FBS-containing media is added. The next day, i.e., 46 hrs post transfection, the media is collected. This media will contain virus particles). Fresh media is added and collected after 24 hours, i.e. 70 hrs post-transfection. The media collected is pooled to process by ultracentrifugation and collect a concentrated titer of virus particles.

### Jmjd3 knockdown in RAW 264.7 macrophages

RAW 264.7 macrophages were transfected with JMJD3 shRNA lentiviral particles for 18-20 h and then incubated for 48 hrs in growth media (+/- statins). Cells were then stimulated with LPS or IL-4 before DNA extraction and qPCR analysis.

### In vivo statin experiment

C57BL/6J mice were fed the chow diet (WQJX Bio-technology) to which simvastatin (100 mg/kg/day; Merck & Co Inc.) added for 14 days. Control animals were fed the chow diet. Peritoneal macrophages were harvested from the mice 4 days after 1.5 mL of thioglycolate broth (Sigma) was injected. Cells were washed with phosphate-buffered saline (PBS), seeded at a density of 1,000,000 per well into 24-well dishes, incubated for 12 hr in Dulbecco’s DMEM containing 10% LPSD and 1 µM simvastatin (to maintain *in vivo* cholesterol levels). Cells were then stimulated with 100 ng/mL LPS (MCE) or 20 ng/mL IL4 (Aladdin), and RNA was isolated for qPCR. Cholesterol contents were analyzed with cholesterol quantification assay kit (Sigma).

### RNA purification and cDNA synthesis

After treatment, the cells were lysed and collected in TRIzol (ThermoFisher Scientific, Watham, MA), and frozen at -80°C. Total RNA was extracted by phenol–chloroform extraction, followed by ethanol precipitation. RNA was purified through columns supplied in the Molecular Biology kit (BioBasic Inc., Markham, ON), according to the manufacturer’s instructions. The RNA concentrations and purity were determined using a NanoDrop One (ThermoFisher Scientific, Watham, MA) spectrophotometer. cDNA synthesis was performed on the Bio-Rad T100 PCR Gradient Thermal Cycler using the QuantiTect® Reverse Transcription kit (Qiagen, Germantown, MD), following manufactureŕs instructions.

### Reverse transcriptase quantitative PCR (RT-qPCR)

Gene expression was analyzed by real time reverse transcriptase quantitative PCR (RT-qPCR) according to the Fast SYBR Green protocol with the AriaMx real-time PCR detection system (Agilent technologies, Santa Clara, CA). Primers were ordered from Invitrogen (ThermoFisher Scientific, Watham, MA) and are listed in Table 1. Each condition was prepared in triplicates and each sample was loaded as technical triplicates for each gene (target or reference) analyzed. The mRNA levels of mouse HPRT1 or GAPDH were used as internal controls (reference gene) as indicated.

**Table 1:**
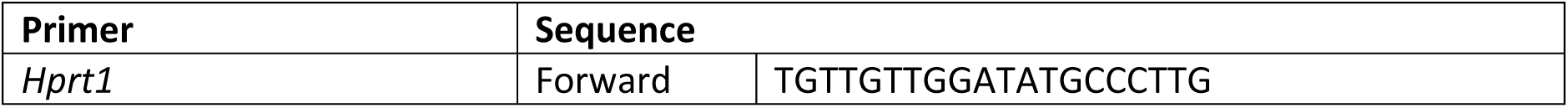
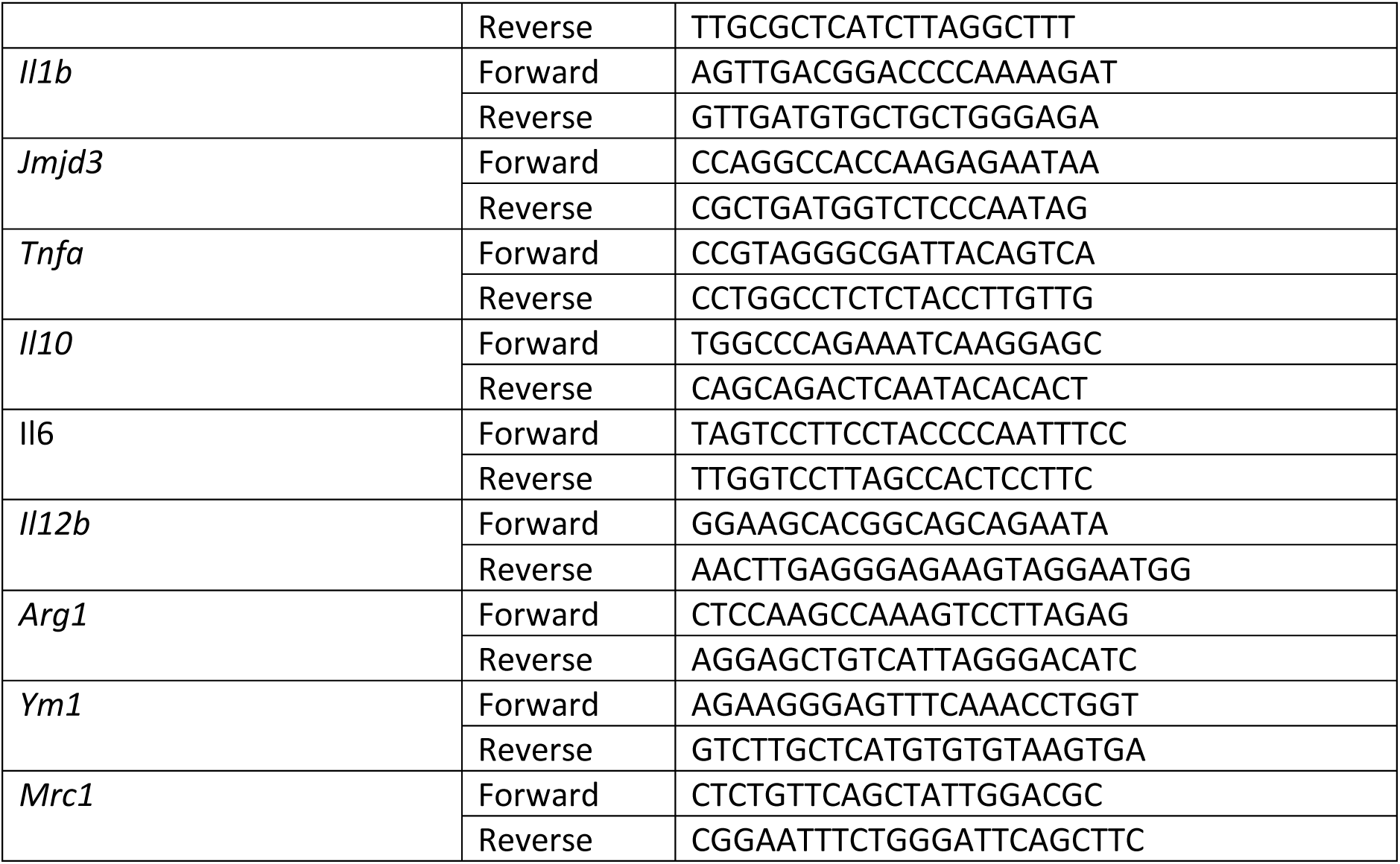
Mouse primers used for real time RT-qPCR.

### Immunoblotting

For Western blot analysis, MCD-treated RAW 264.7 cells were washed in ice-cold PBS, lysed in radioimmune precipitation assay (RIPA) buffer (150 mM NaCl, 1% Nonidet P-40, 1% sodium deoxycholate, and 25 mM Tris (pH 7.6)) supplemented with a cocktail of protease and phosphatase inhibitors. Total cell lysis was achieved through sonication for 20 seconds at 20% amplitude and samples were stored at -80°C. Histone extraction was performed following a specific protocol from Abcam. Protein concentration was determined using the Bradford assay (Bio-Rad, 5000006). SDS buffer (313 mM Tris (pH 6.8), 10% SDS, 0.05% bromophenol blue, 50% glycerol, and 0.1 M DTT) was added, and the samples were boiled at 100 °C for 5 minutes. Proteins were separated by SDS-PAGE on 10% acrylamide gels. After separation, proteins were transferred onto PVDF membranes and were blocked in 5% milk powder (in PBS, 1% Triton X-100) for 1 h. Membranes were incubated with the primary antibodies in the following conditions: overnight incubation with the anti-Jmjd3 antibody (1:400) at 4°C, and 1h with the anti-H3K27Me3 antibody at room temperature. After washes, blots were further incubated with an HRP-conjugated anti-rabbit antibody (1:10 000) for 1h. Blots were imaged using ECL-based film detection system.

### MCD/cholesterol (10:1 mol/mol) preparation

Cholesterol (0.3 mmol) of is dissolved in 1 ml chloroform solution and dried under nitrogen in a glass culture tube. 10 ml of a MCD solution (300 mM) prepared in BSA/BSS buffer solution (20 mM Hepes, pH 7.4, 135 mM NaCl, 5 mM KCl, 1.8 mM CaCl_2_, 1 mM MgCl_2_, 5.6 mM glucose, and 1 mg/ml BSA) was added to the tube, and the resulting suspension was vortexed, and bath-sonicated at 37°C until the suspension clarified. The suspension was then incubated in a rocking water bath overnight at 37°C to maximize formation of soluble complexes. This is the stock solution that can be diluted into desired concentrations.

### Macrophage cholesterol depletion and repletion

On the day of experiment, BMDMs are incubated with 5 mM MCD for 1 h. The cells are rinsed in 0.25% BSA in DMEM and left to rest for 10 minutes before switching the media to media containing 1 mM MCD/cholesterol complex for 1 h. RNAs are collected at the end of incubation for *Jmjd3* expression. Controls cells will be cells that did receive any treatment or MCD. Triplicates wells are prepared and Jmjd3 expression was determined by RT-PCR.

### NF-kB activation assay

NF-kB activation was determined by performing the QUANTI-Blue assay using the RAW-Blue^TM^ cells (InvivoGen). Through exposure of the RAW-Blue cells to various substances, NF-kB/AP-1 activation is induced, making secreted embryonic alkaline phosphatase (SEAP) into the cell supernatant. In brief, RAW-Blue cells were seeded in 6-well plates (1.5×10^5^ cells/ml) and were allowed to recover for 2 days in growth media (similar to RAW264.7 cells) before they were stimulated with LPS (100 ng/ml) or MCD (5 mM) for 3h. Then, the cells and medium for each sample were collected and sonicated on ice for 15 seconds at 20% amplitude. The supernatant was separated from cell debris by centrifugation at 4°C for 10 minutes at 10,000 rpm. For detection, the cell supernatant and a secreted embryonic alkaline phosphatase (SEAP) detection reagent (QUANTI-Blue) were mixed and the absorbance was measured at 650 nm as described in the assay instructions. Each condition was performed in triplicate samples and each sample was measured in technical triplicates.

### Mitochondrial Oxygen Consumption

Cellular oxygen consumption rates (OCR) were measured using an extracellular flux analyzer (Seahorse XF96e; Agilent Technologies, Santa Clara, CA). Briefly, the cartridge sensors were incubated overnight at 37 °C in a hydration/ calibration solution (XF Calibrant; Agilent Technologies). Specially designed polystyrene tissue culture-treated 96-well microplates with a clear flat bottom (Seahorse XF96 V3 PS Cell Culture Microplates; Agilent Technologies) were then seeded with 80 µl of cell suspension (4.4×10^5^ cells/ml) per well and incubated 2–3 h under cell culture conditions to allow cell attachment. For treatment with MCD, cells were washed with pre-warmed sterile PBS and the culture supernatants were replaced with medium containing MCD (5 mM). Then, the cells were incubated for 1 h under cell culture conditions. For treatment with statins, cells were washed with pre-warmed sterile PBS and the culture supernatants were replaced with medium supplemented with lipoprotein-deficient serum (LPDS), containing statins (7 mM lovastatin + 200 mM mevalonate). Then, the cells were incubated for 2 days under cell culture conditions. At the end of either treatment, the cells are washed and incubated 45 minutes at 37 °C in extracellular flux analysis medium (sodium bicarbonate-, glucose-, phenol red-, pyruvate-, and glutamine-free modified DMEM [Sigma-Aldrich, catalog no. D5030] freshly supplemented with cell culture-grade D-glucose (4.5 g/L), cell culture-grade L-glutamine [4 mM; Wisent], and 4-(2-hydroxyethyl)-1-piperazineethanesulfonic acid [HEPES; 4.5 mM; Sigma-Aldrich], pH adjusted to 7.35-7.40 at room temperature). OCR were measured to assess resting respiration, followed by ATP production-dependent respiration, maximal respiration, and non-mitochondrial oxygen consumption, after sequential injections of oligomycin (an ATP synthase inhibitor) at 1µM (final concentration), CCCP (an ionophore acting as a proton uncoupler) at 2µM, and rotenone together with antimycin A (complex I and complex III inhibitors, respectively), each at 0.5µM. mitochondrial ATP production-dependent respiration was calculated by subtracting the lowest OCR after oligomycin injection from resting OCR. For each parameter, OCR measurements were performed at least three times at 6-minute intervals. Each condition was performed in 7–8 replicates samples. All OCR measurements were corrected for the OCR of cell-free wells containing only medium. Upon completion of the OCR measurements, the cells were washed once with PBS and lysed in 1M NaOH (40 µl/well). The lysates were kept at 4 °C for up to 24 h, and protein determination was performed using the Bradford colorimetric assay with BSA as the standard protein (Thermo Scientific). Absorbance was measured at a wavelength of 595nm using a hybrid microplate reader (Biotek).

### 27-Hydroxycholesterol (27-HC) quantification

Cell pellets were collected from RAW264.7 macrophages treated with 1, 3 and 5 mM of MCD for 1 hour. Samples were sent to the University of Texas (UT) Southwestern Medical center for analysis by ultraperformance liquid chromatography/electrospray ionization/tandem mass spectrometry allowing chromatographic resolution of the hydroxycholesterol species. 27-HC levels were calculated by normalizing the 27-HC amounts to the protein content in the whole cell pellet.

### Purification and formation of E. coli inner membrane vesicles (IMVs)

*E. coli* inner membranes were purified from *E. coli* MG1655 strain by sucrose gradient ultracentrifugation (30 minutes at 80,000 rpm, Optima^TM^ MAX-XP) as previously described^2^. The *E.coli* inner membranes were then resuspended in either the synthesis buffer (MES 100 mM pH 6 and NaCl 25 mM) or the hydrolysis buffer (HEPES 25 mM pH 8) and ultrasonicated for 15 minutes (5 seconds on-off cycles at 30% power, VCX500 Ultrasonic Processors). The resulting homogeneous solution contained the inner membrane vesicles (IMVs).

### Loading and depletion of cholesterol in IMVs

Prior to any treatment, the protein concentration of IMVs was determined by the BCA protein assay and IMVs were diluted to a final protein concentration of 0.5 mg/ml. IMVs were then loaded with cholesterol using a modified protocol based on the MCD/cholesterol complex^3^. To load IMVs, 25 mM MCD were mixed with 2.5 mg/ml of cholesterol on HEPES (25 mM, pH7.8) following the protocol described in^4^. After incubation with MCD-cholesterol complexes, IMVs were ultracentrifuged (80,000 rpm for 30 minutes) and the supernatant removed. The IMVs were then resuspended in HEPES 25 mM (pH 7.8). After loading, the lipid, cholesterol and protein concentration of IMVs were determined by the phosphate determination assay^5^, Amplex Red cholesterol assay kit^6^ and BCA protein assay^7^, respectively. To achieve vesicles with different cholesterol content, cholesterol-loaded IMVs (1mg/mL protein concentration) were treated with increasing MCD concentrations (from 0 to 7.0 mM) for 30 minutes at 37°C^3, 8^. Cholesterol-depleted vesicles were ultracentrifuged (80000 rpm for 30 minutes) and the pellet was resuspended in HEPES (25 mM, pH=7.8). Lipid, protein and cholesterol concentration of depleted samples were quantified again after the MCD treatment.

### Hydrolysis and synthesis of ATP on cholesterol doped IMVs

ATP hydrolysis was performed by adding a total concentration of 2 mM ATP to 200 mM IMVs (lipid concentration) and incubated for 30 minutes. The concentration of phosphates from ATP hydrolysis was measured using the malaquite green assay^9^. ATP synthesis was triggered by promoting a ΔpH across the IMV membranes by mixing the samples (IMVs resuspended in MES 100 mM pH 6) with an external buffer with higher pH and in the presence of inorganic phosphate, ADP and magnesium ion (HEPES 100 mM pH 8, 5 mM P_i_, 2.5 mM ADP, 5 mM MgCl_2_, 25 mM NaCl) at a volume ratio of 1:10 and incubated for 2 to 5 minutes. The reaction was stopped by adding 20% TCA at a ratio volume of 10:1, and then the samples was equilibrated to neutral pH^10^. ATP concentration after synthesis was measured using ATP detection assay kit (Molecular Probes) with a luminometer GloMax®-Multi Detection. Amplex™ Red Cholesterol Assay Kit and Pierce® BCA Protein assay kits were supplied by ThermoFisher. Luminescent ATP Detection Assay Kit (based on firefly’s luciferase / luciferin) was purchased from Molecular Probes.

### RNA-seq and data analysis

RAW 264.7 macrophages in duplicates were either left untreated or treated with 5 mM MCD for 1 hour. Total RNA was purified as described above, and the RNA concentrations and purity were determined using a NanoDrop One (ThermoFisher Scientific, Watham, MA) spectrophotometer. Library preparation and 150-bp paired-end RNA-Seq were performed using standard Illumina procedures for the NextSeq 500 platform. Reads libraries produced from RNA sequencing for each replicate and condition were aligned against mouse genome reference provided by the GENCODE project – the mouse genome reference release M25^11^. Transcript quantification count tables were generated per bait screening using the Salmon algorithm ver1.7.0^12^. The following comparisons: MCD1 vs Control, Statin vs Control, and Statin-LPS vs LPS were performed to identify differentially expressed (DEGs) genes using DESeq2 ver1.40.2^13^.

### GO enrichment and KEGG pathway analysis

Gene ontology functional enrichment (GO) analysis and KEGG pathway analysis for the DEGs was performed using clusterProfiler ver4.8.2^14^ to identify significantly enriched biological processes associated with the set of genes upregulated in the experimental condition greater than log2 fold change of 1 and with the set of genes downregulated with log2 fold change of less than -1. A p value cut off of 0.05 was defined for the analyses. The Benjamini-Hochberg method was applied to adjust the p-values for multiple testing. This method controls the false discovery rate during the adjustment process.

### ATAC-seq

RAW 264.7 macrophages in duplicates were either left untreated or treated with 5 mM MCD for 1 hour. Cells were detached using 0.5% trypsin and resuspended in chilled 1X PBS containing 1 mM EDTA. Visible cell numbers per sample were obtained by staining the cells with trypan blue and counting them using hemocytometer. 50,000 cells were used for performing ATAC-Seq. Cells were washed in 100 µl of ice cold 1X PBS and centrifuged at 500 x g for 5 minutes. Nuclei are prepared by lysing the cells in ice cold lysis buffer containing 10 mM Tris-Cl, pH7.4, 10 mM NaCl, 3 mM MgCl_2_ and 0.1% IGEPAL CA-630. Nuclei were pelleted by spinning the samples at 500 x g for 10 minutes in fixed-angle cold centrifuge and proceeded for tagmentation reaction. A 25 µl tagmentation reaction was setup by resuspending the nuclei in 12.5 µl of 2x Tagment DNA buffer and 5 µl of TDE1 transposase. Samples were incubated at 37°C for 30 minutes with gentle intermittent mixing. Following transposition, the sample volume was made up to 50 µl using resuspension buffer and were processed for DNA preparation. For tagmented DNA clean-up, 180 µl of Zymo DNA binding buffer was added and mixed thoroughly before loading on to Zymo-spin concentrator-5 columns. For transposase free DNA, the samples were eluted in 16.5 µl of elution buffer. Purified tagmented DNA was amplified with KAPA HIFI polymerase (12 PCR cycles) and Unique Dual Primers from Illumina (Cat number 20332088). Size-selection (L:1.1; R :0.6) was performed with KAPA Pure beads and size distribution of the final libraries was assessed on bioanalyzer (Agilent). Libraries were quantified by qPCR and loaded equimolarly on a S4 Novaseq flowcell. Each library was sequenced with a coverage of 50M paired-end reads (PE100).

### ATAC-seq analysis

Sequencing reads for chromatin accessibility (ATAC) were aligned to mus musculus genome assembly GRCm38 (mm10) using Bowtie2^15^ with default parameters. The resulting BAM files were filtered to remove duplicate reads using Picard Tools (https://broadinstitute.github.io/picard/). Peaks were called using MACS2 ver2.0^16^ (Zhang et al, 2008) with the parameter “-nomodel”. The generated narrow peaks files were used for downstream analysis. Diffbind ver3.10^17,18^ was used to identify differentially accessible regions (peaks) called by MACS2. Comparison between the 2 replicates of the 2 conditions MCD1 vs Control was conducted. DESeq2 ver1.40.2^19^ and EdgeR ver3.42.4^20^ were used within Diffbind to identify regions of differential accessible between MCD1 treated and control (p value <0.05 and RD <0.05). ChipSeeker ver1.3 ^21,22^ was used to annotate the genomic features of the differentially accessible peaks identified by Diffbind, where the maximum range of promoter to transcription start site was set to 3kb. The peaks were assigned to the nearest genes based on distance of the peak region to the transcription start site. This allowed the annotation of ATAC-seq peaks with genes. The heatmap was generated using Diffbind::dba.plotProfile in order to compute peakset profiles for MCD1 and control conditions of loss or gain of genomic accessibility.

### Gene Set Enrichment Analysis

Functional Gene Set Enrichment Analysis (FGSEA) was conducted for all RNA-seq comparisons and ATAC-seq comparison. FGSEA was conducted using fgsea ver1.26^23^. MSigDB gene sets utilized in fgsea were Hallmark gene sets ver7.1, C2 BioCarta pathways ver7.1, C2 KEGG pathways ver7.1, C2 Reactome pathways ver7.1, and C3 Transcription factor targets ver7.1. The gene list provided to fgsea were based on the DEGs, including both upregulated and downregulated genes ranked in descending order or log2 fold change. The size of the gene sets considered for the enrichment analysis was set to a minimum of 15 and maximum of 500. Only Hallmark pathways with adjusted p value of less than 0.01 was plotted. Enrichment plots for the Hallmark pathway with the highest normalized enrichment score (NES) are illustrated.

### Statistical analysis

Statistical analyses between data groups were performed with PRISM software (GraphPad). Data for real time RT-qPCR and Seahorse experiments are presented as the mean ± S.D. as indicated. The statistical significance of differences between groups was analyzed by Student’s *t* test. Differences were considered significant at a *p* value < 0.05.

---

**Figure 1 - figure supplement 1.**
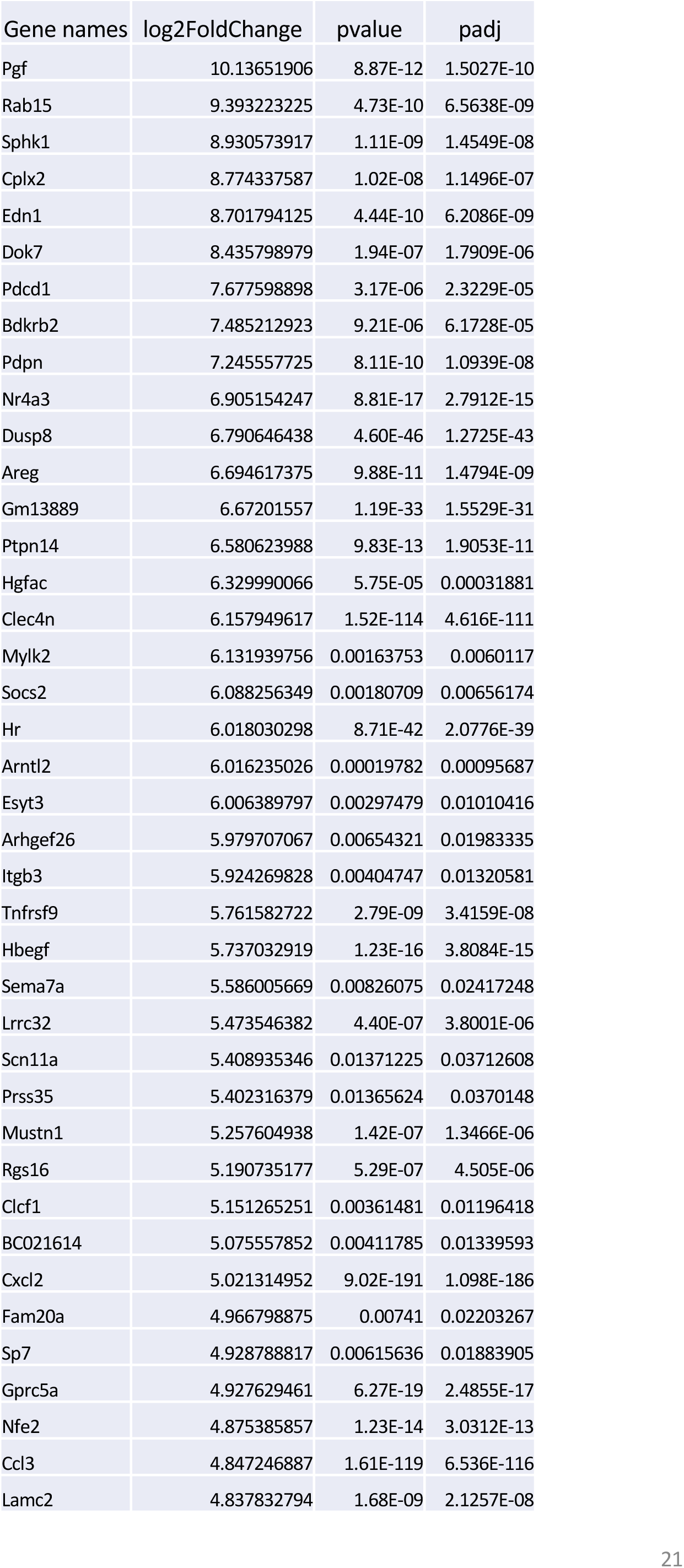
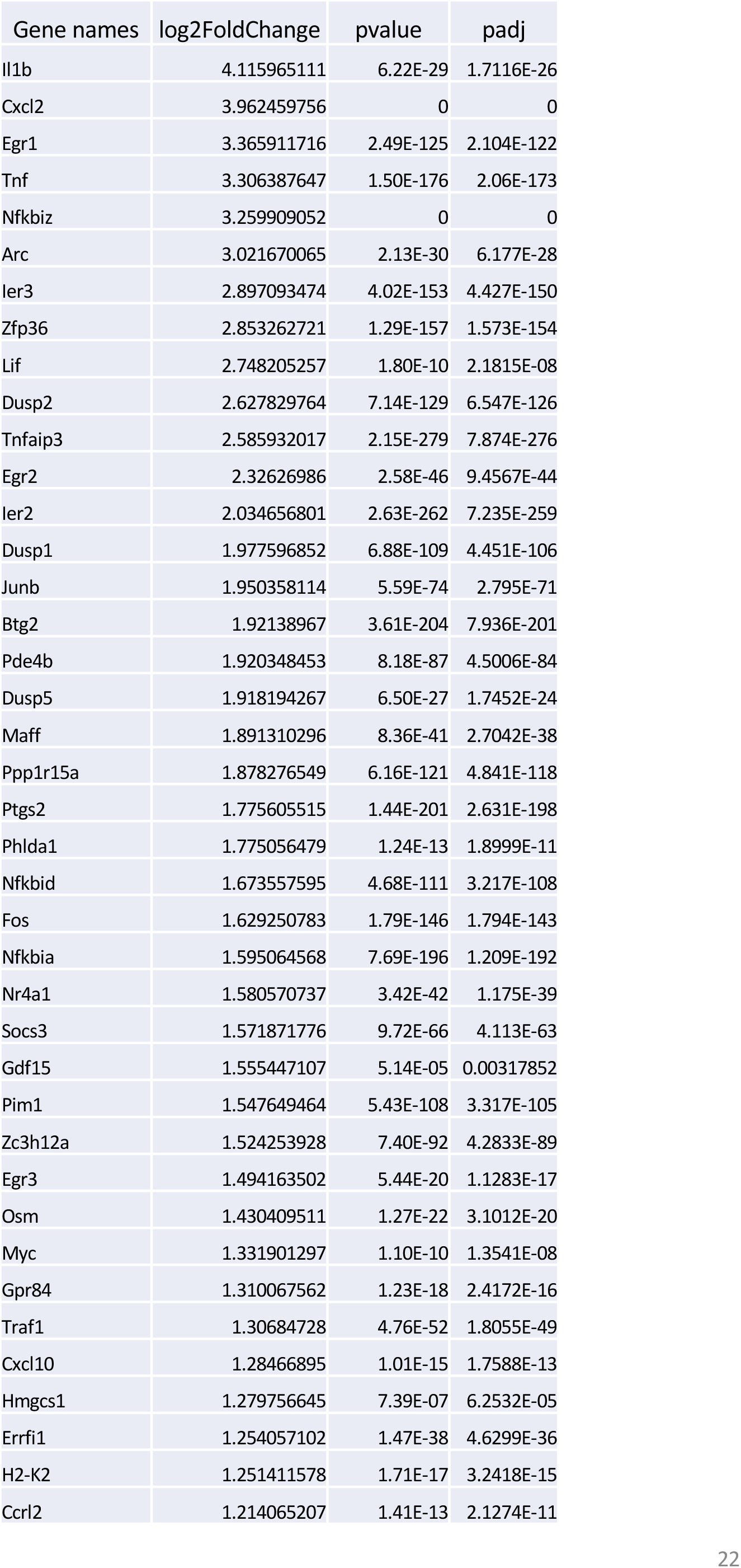
Upregulated genes (top 40) by statin and MCD. RNA-seq was performed in statin- or MCD-treated macrophages. For genes upregulated (log2>1) (Supplementary file 1), the top 40 upregulated genes by statin **(A)** and by MCD **(B).**

**Figure 1 - figure supplement 2.**
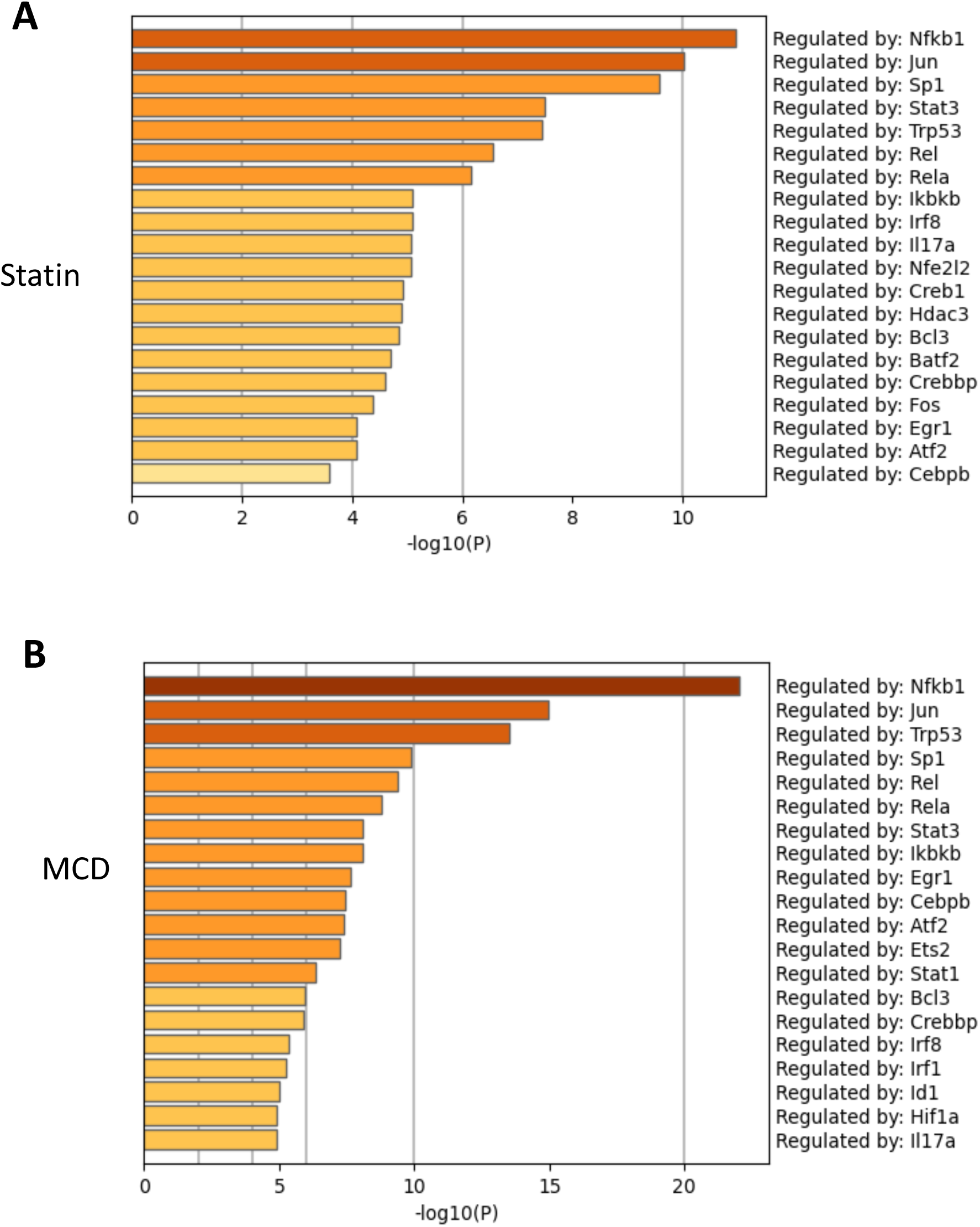
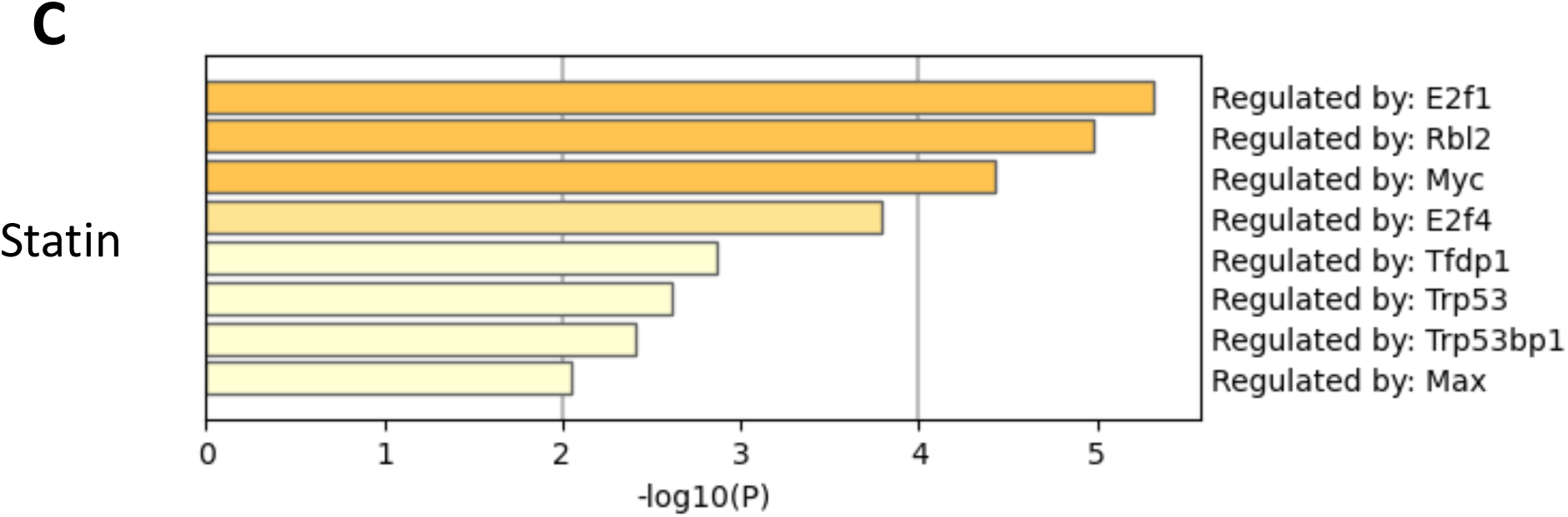
Cellular cholesterol contents regulate NF-kB pathway. RAW macrophages were treated with MCD or statin and RNA-seq was performed as in Figure. 1. TF regulatory pathways of activated genes were analyzed using Metascape; top TFs employed by either lovastatin (***A***) (7 µM + 200 µM mevalonate); or MCD *(**B**)* (5mM, 1h). The genes down-regulated by statin is analyzed in (**C**). No TF is identified in MCD down-regulated genes.

**Figure 1 - figure supplement 3.**
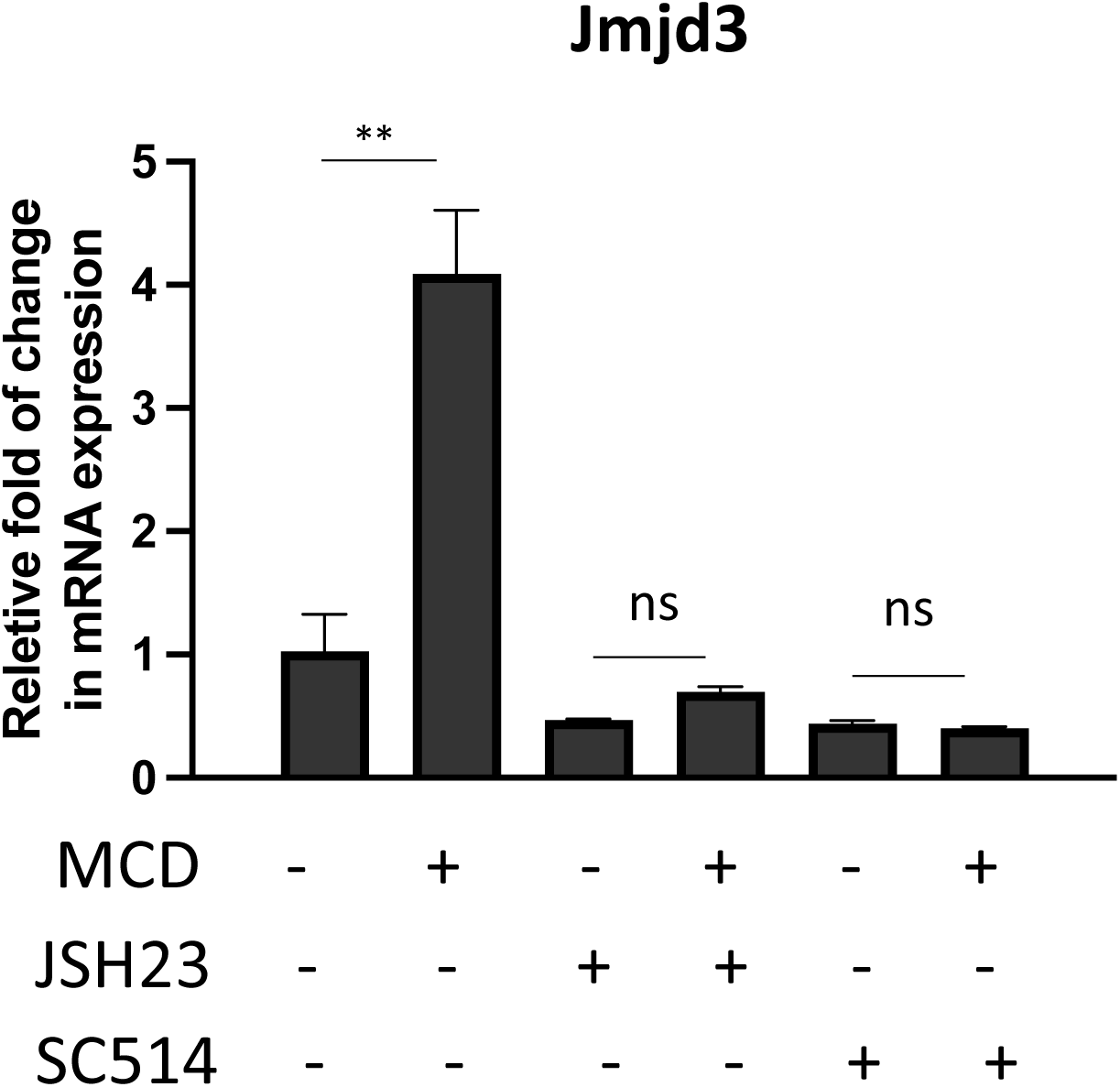
The expression of *Jmjd3* activated by MCD requires NF-κB activity. Effect of NF-kB inhibitors, JSH23 (10 µM) and SC514 (10 µM), on *Jmjd3* expression in MCD-treated RAW macrophages. Graphs are representative of 3 independent experiments with 3 replicates per condition and are presented as means ± SD. Statistical analysis was performed using unpaired, two-tailed Student’s t-test. An asterisk (*) or double asterisks (**) indicate a significant difference with p<0.05 and p<0.001, respectively.

**Figure 1 - figure supplement 4.**
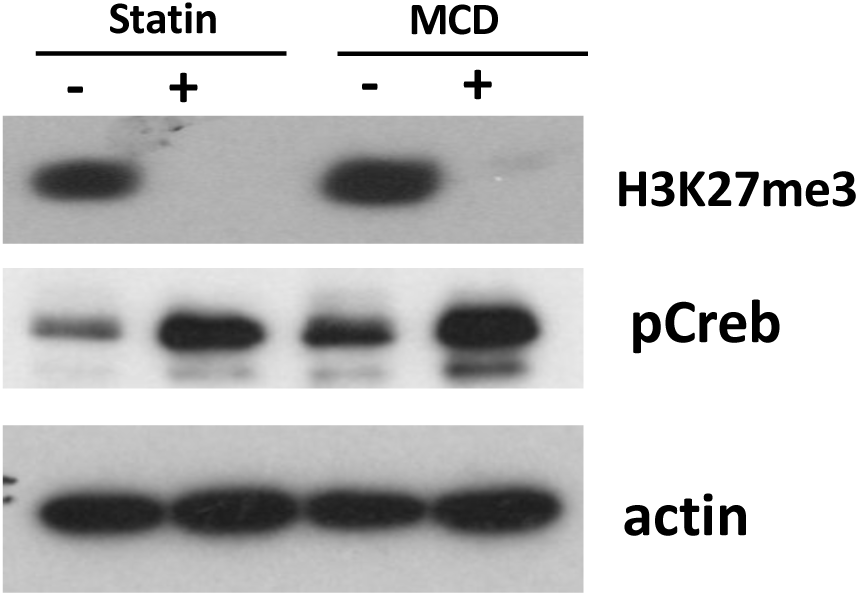
Levels of H3K27Me3 are decreased in macrophages treated with statin or MCD. (lovastatin, 7 µM + 200 µM mevalonate; 2 days) or MCD (5mM, 1h). The pCREB was used as internal control for cholesterol depletion and actin a loading control. Original blots are in source data.

**Figure 4 - figure supplement 5.**
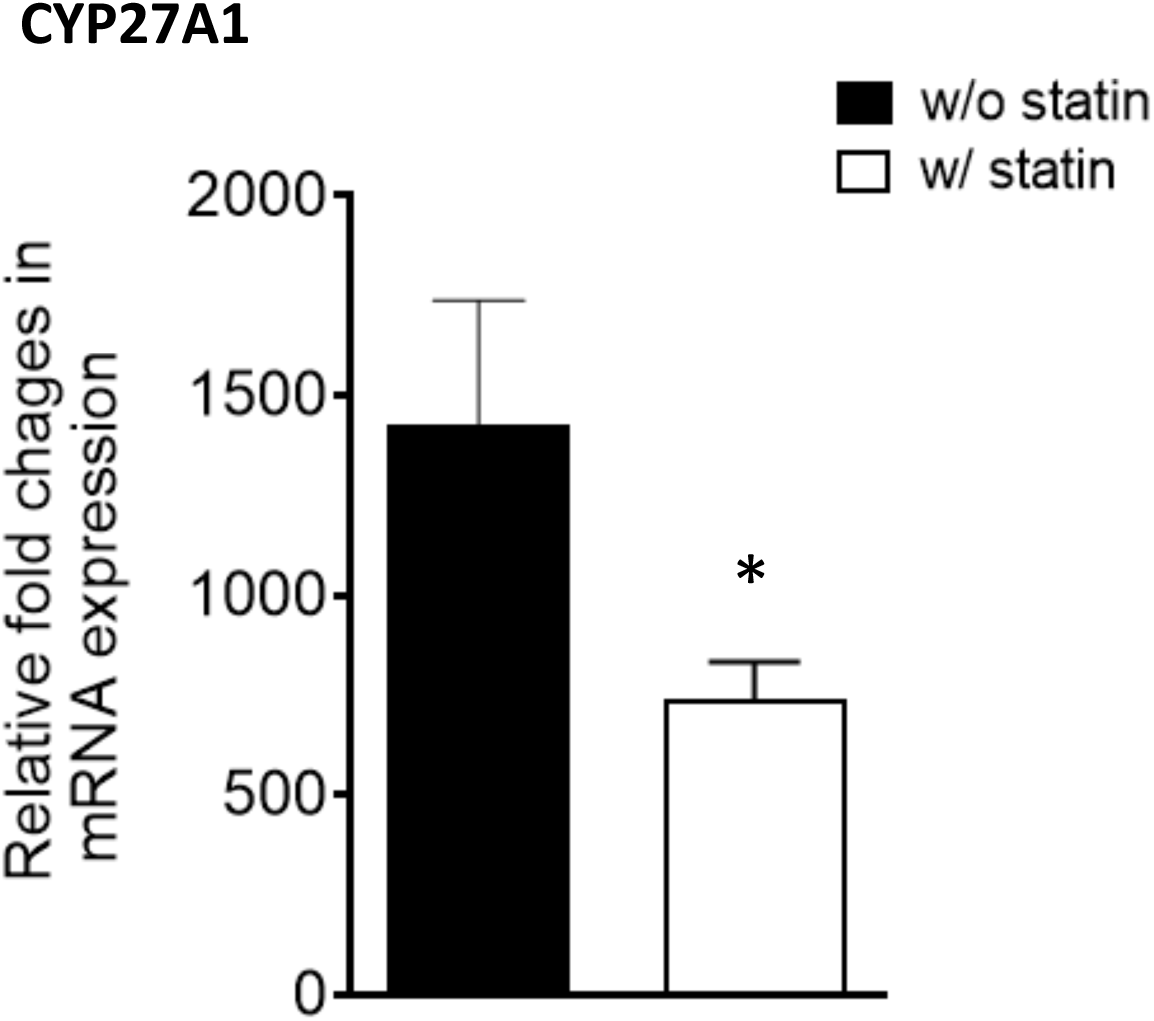
Sterol 27-hydroxylase, *Cyp27a1*, expression. RAW 264.7 macrophages were treated with compactin (7 µM + 200 µM mevalonate) for 2 d and the expression of *Cyp27a1* analyzed. P<0.01.

**Figure 5 - figure supplement 6.**
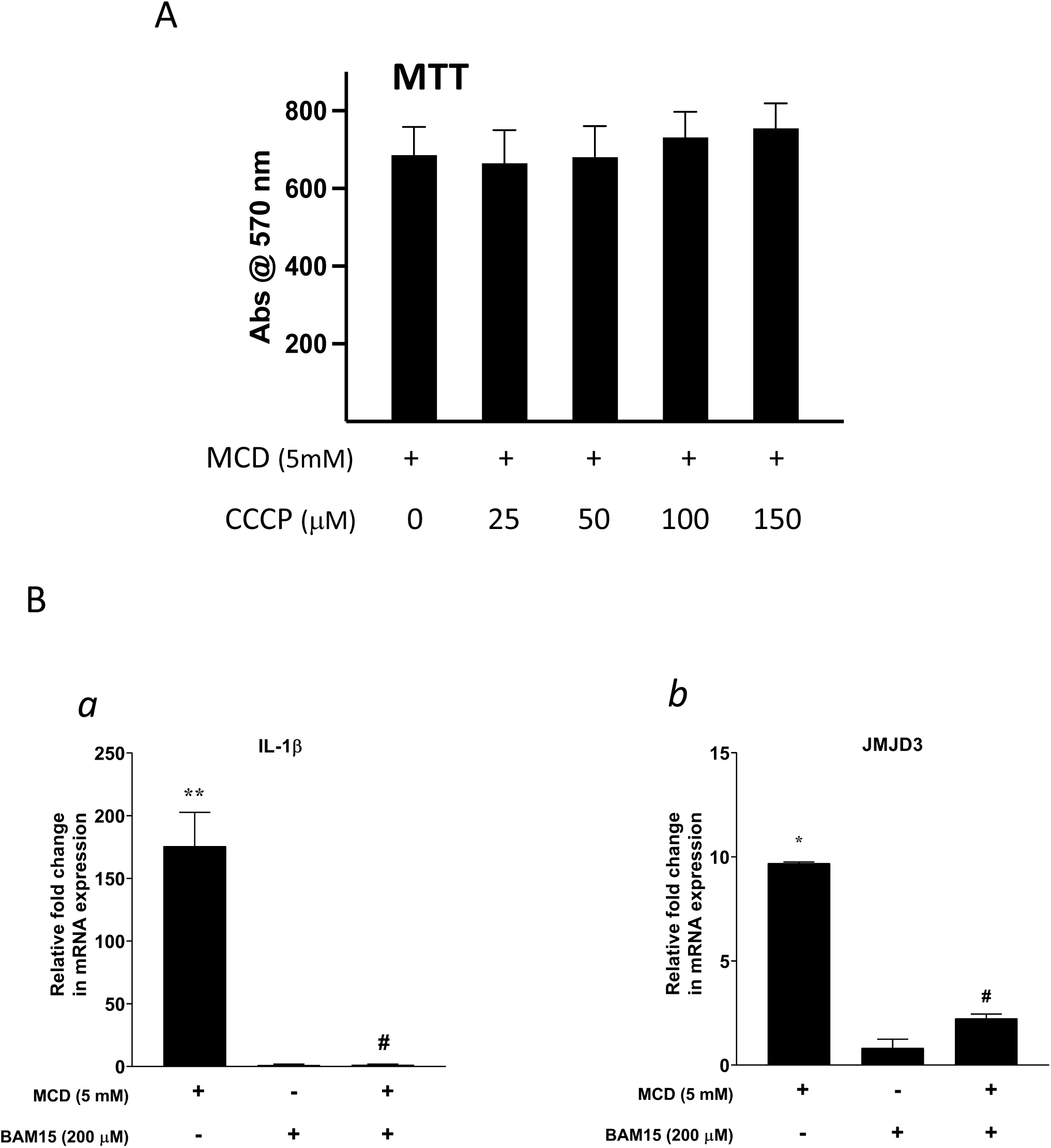
The effect of proton flux Inhibitors. **(A)** The toxicity of CCCP by MTT assay. **(B)** Effect of BAM15 (200 µM) on *Il1b (**a**)* and *Jmjd3 (**b***) gene expression in MCD-treated RAW264.7 macrophages. Data is representative of 3 independent samples per condition and are presented as mean ± SD. Statistical analysis was performed using unpaired, two-tailed Student’s t-test. asterisk (*) and hashtag (#) indicate a significant difference between MCD without or with inhibitors with p < 0.05.

**Figure 6 - figure supplement 7.**
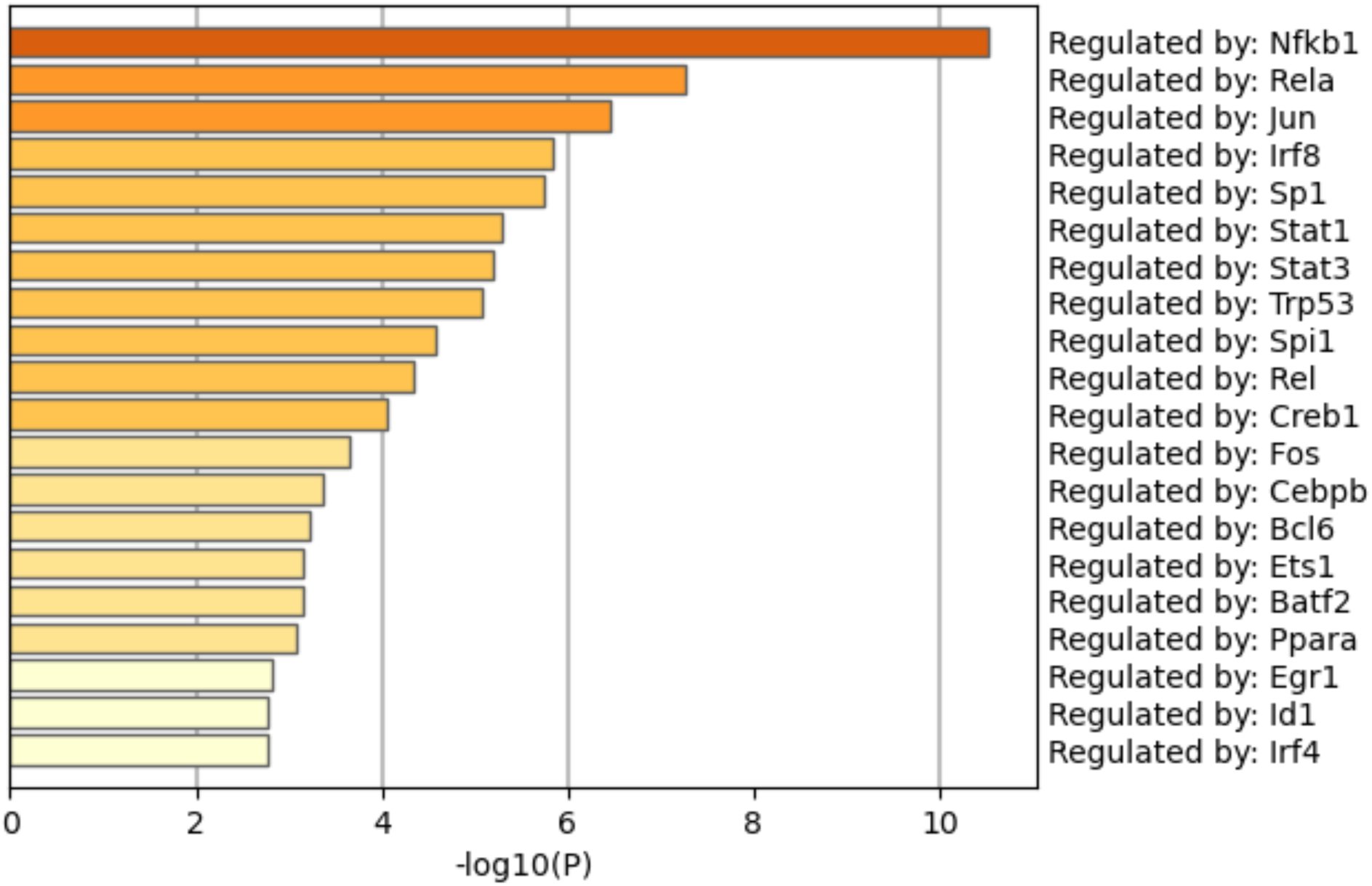
Pathway analysis among the genes opened by MCD in ATAC-seq: Regulatory pathways of genes opened by MCD were analyzed using Metascape.

**Figure 8 - figure supplement 8.**
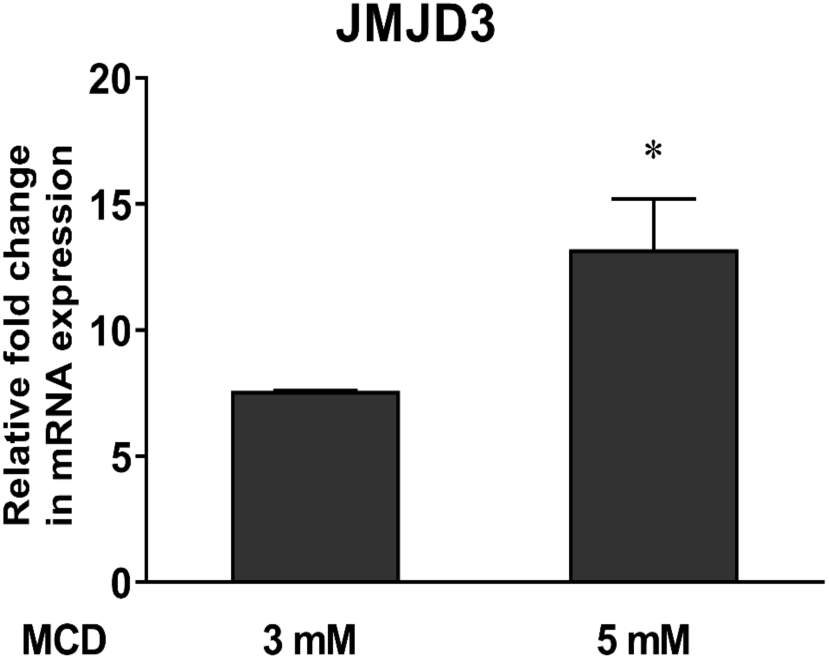
Expression of the Jmjd3 in 264.7 macrophages with increasing doses of MCD (mean ± SD). An asterisk (*) indicates a significant difference with p<0.01.

**Figure 8 - figure supplement 9.**
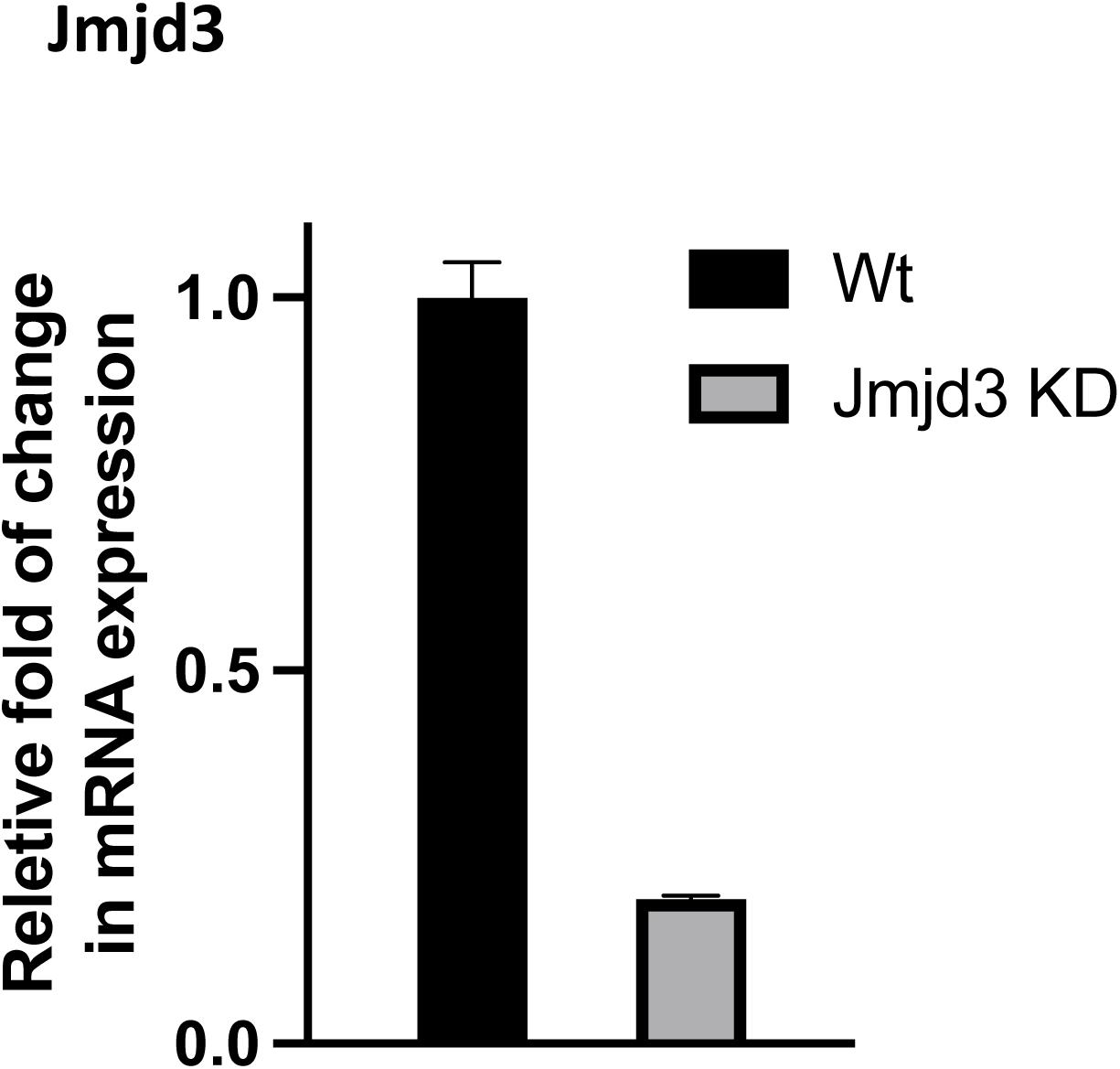
Expression of JMJD3 in wt and *Jmjd3* KD macrophages (mean ± SD), P<0.001.

## Footnotes

1 Lovastatin, compactin, simvastatin and pravastatin were used exchangeably throughout the study without significant differences in results.

## References

1. Tabas I, Glass CK. Anti-inflammatory therapy in chronic disease: challenges and opportunities. Science. 2013 Jan 11;339(6116):166–72.

2. Tabas I, Lichtman AH. Monocyte-Macrophages and T Cells in Atherosclerosis. Immunity. 2017 Oct 17;47(4):621–634.

3. Levy Y, Leibowitz R, Aviram M, et al. Reduction of plasma cholesterol by lovastatin normalizes erythrocyte membrane fluidity in patients with severe hypercholesterolaemia. Br J Clin Pharmacol 1992; 34 (5): 427–30

4. Hochgraf E, Levy Y, Aviram M, et al. Lovastatin decreases plasma and platelet cholesterol levels and normalizes elevated platelet fluidity and aggregation in hypercholesterolemic patients. Metabolism 1994; 43 (1): 11–7

5. Lijnen P, Celis H, Fagard R, et al. Influence of cholesterol lowering on plasma membrane lipids and cationic transport systems. J-Hypertens 1994; 12 (1): 59–64.

6. Parihar SP, Guler R, Khutlang R, Lang DM, Hurdayal R, Mhlanga MM, Suzuki H, Marais AD, Brombacher F. Statin therapy reduces the mycobacterium tuberculosis burden in human macrophages and in mice by enhancing autophagy and phagosome maturation. J Infect Dis. 2014 Mar 1;209(5):754–63.

7. Libby P. Inflammation in Atherosclerosis-No Longer a Theory. Clin Chem. 2021 Jan 8;67(1):131–142.

8. Ma L, Dong F, Zaid M, Kumar A, Zha X. ABCA1 protein enhances Toll-like receptor 4 (TLR4)-stimulated interleukin-10 (IL-10) secretion through protein kinase A (PKA) activation. J Biol Chem. 2012 Nov 23;287(48):40502–12.

9. Yvan-Charvet L, Welch C, Pagler TA, Ranalletta M, Lamkanfi M, Han S, Ishibashi M, Li R, Wang N, Tall AR. Increased inflammatory gene expression in ABC transporter-deficient macrophages: free cholesterol accumulation, increased signaling via toll-like receptors, and neutrophil infiltration of atherosclerotic lesions. Circulation. 2008 Oct 28;118(18):1837–47.

10. Zhu X, Lee JY, Timmins JM, Brown JM, Boudyguina E, Mulya A, Gebre AK, Willingham MC, Hiltbold EM, Mishra N, Maeda N, Parks JS. Increased cellular free cholesterol in macrophage-specific Abca1 knock-out mice enhances pro-inflammatory response of macrophages. J Biol Chem. 2008 Aug 22;283(34):22930–41.

11. Raetz CR, Garrett TA, Reynolds CM, Shaw WA, Moore JD, Smith DC Jr, Ribeiro AA, Murphy RC, Ulevitch RJ, Fearns C, Reichart D, Glass CK, Benner C, Subramaniam S, Harkewicz R, Bowers-Gentry RC, Buczynski MW, Cooper JA, Deems RA, Dennis EA. Kdo2-Lipid A of Escherichia coli, a defined endotoxin that activates macrophages via TLR-4. J Lipid Res. 2006 May;47(5):1097–111.

12. Link VM, Gosselin D, Glass CK. Mechanisms Underlying the Selection and Function of Macrophage-Specific Enhancers. Cold Spring Harb Symp Quant Biol. 2015; 80:213–21.

13. Troutman TD, Kofman E, Glass CK. Exploiting dynamic enhancer landscapes to decode macrophage and microglia phenotypes in health and disease. Mol Cell. 2021 Oct 7;81(19):3888–3903.

14. Spann NJ, Garmire LX, McDonald JG, Myers DS, Milne SB, Shibata N, Reichart D, Fox JN, Shaked I, Heudobler D, Raetz CR, Wang EW, Kelly SL, Sullards MC, Murphy RC, Merrill AH Jr, Brown HA, Dennis EA, Li AC, Ley K, Tsimikas S, Fahy E, Subramaniam S, Quehenberger O, Russell DW, Glass CK. Regulated accumulation of desmosterol integrates macrophage lipid metabolism and inflammatory responses. Cell. 2012 Sep 28;151(1):138–52

15. Ivashkiv LB. Epigenetic regulation of macrophage polarization and function. Trends Immunol. 2013 May;34(5):216–23.

16. Mayor S, Sabharanjak S, Maxfield FR. Cholesterol-dependent retention of GPI-anchored proteins in endosomes. EMBO J. 1998 Aug 17;17(16):4626–38.

17. Han H, Cho JW, Lee S, Yun A, Kim H, Bae D, Yang S, Kim CY, Lee M, Kim E, Lee S, Kang B, Jeong D, Kim Y, Jeon HN, Jung H, Nam S, Chung M, Kim JH, Lee I. TRRUST v2: an expanded reference database of human and mouse transcriptional regulatory interactions. Nucleic Acids Res. 2018 Jan 4;46(D1):D380–D386.

18. Zhou Y, Zhou B, Pache L, Chang M, Khodabakhshi AH, Tanaseichuk O, Benner C, Chanda SK. Metascape provides a biologist-oriented resource for the analysis of systems-level datasets. Nat Commun. 2019 Apr 3;10(1):1523.

19. De Santa F, Totaro MG, Prosperini E, Notarbartolo S, Testa G, Natoli G. The histone H3 lysine-27 demethylase Jmjd3 links inflammation to inhibition of polycomb-mediated gene silencing. Cell. 2007 Sep 21;130(6):1083–94

20. De Santa F, Narang V, Yap ZH, Tusi BK, Burgold T, Austenaa L, Bucci G, Caganova M, Notarbartolo S, Casola S, Testa G, Sung WK, Wei CL, Natoli G. Jmjd3 contributes to the control of gene expression in LPS-activated macrophages. EMBO J. 2009 Nov 4;28(21):3341–52.

21. Hansen FC, Kalle-Brune M, van der Plas MJ, Strömdahl AC, Malmsten M, Mörgelin M, Schmidtchen A. The Thrombin-Derived Host Defense Peptide GKY25 Inhibits Endotoxin-Induced Responses through Interactions with Lipopolysaccharide and Macrophages/Monocytes. J Immunol. 2015 Jun 1;194(11):5397–406.

22. Tannahill GM, Curtis AM, Adamik J, Palsson-McDermott EM, McGettrick AF, Goel G, Frezza C, Bernard NJ, Kelly B, Foley NH, Zheng L, Gardet A, Tong Z, Jany SS, Corr SC, Haneklaus M, Caffrey BE, Pierce K, Walmsley S, Beasley FC, Cummins E, Nizet V, Whyte M, Taylor CT, Lin H, Masters SL, Gottlieb E, Kelly VP, Clish C, Auron PE, Xavier RJ, O’Neill LA. Succinate is an inflammatory signal that induces IL-1β through HIF-1α. Nature. 2013 Apr 11;496(7444):238–42.

23. Jha AK, Huang SC, Sergushichev A, Lampropoulou V, Ivanova Y, Loginicheva E, Chmielewski K, Stewart KM, Ashall J, Everts B, Pearce EJ, Driggers EM, Artyomov MN. Network integration of parallel metabolic and transcriptional data reveals metabolic modules that regulate macrophage polarization. Immunity. 2015 Mar 17;42(3):419–30.

24. Steck TL, Lange Y. Cell cholesterol homeostasis: mediation by active cholesterol. Trends Cell Biol. 2010 Nov;20(11):680–7. doi: 10.1016/j.tcb.2010.08.007. Epub 2010 Sep 16. PMID: 20843692; PMCID: PMC2967630.

25. Lange Y, Steck TL. Active cholesterol 20 years on. Traffic. 2020 Nov;21(11):662–674.

26. Lange Y, Steck TL, Ye J, Lanier MH, Molugu V, Ory D. Regulation of fibroblast mitochondrial 27-hydroxycholesterol production by active plasma membrane cholesterol. J Lipid Res. 2009 Sep;50(9):1881–8.

27. Li X, Hylemon P, Pandak WM, Ren S. Enzyme activity assay for cholesterol 27-hydroxylase in mitochondria. J Lipid Res. 2006 Jul;47(7):1507–12.

28. van der Does C, de Keyzer J, van der Laan M, Driessen AJ. Reconstitution of purified bacterial preprotein translocase in liposomes. Methods Enzymol. 2003;372:86–98.

29. Meyer A, Laverny G, Bernardi L, Charles AL, Alsaleh G, Pottecher J, Sibilia J, Geny B. Mitochondria: An Organelle of Bacterial Origin Controlling Inflammation. Front Immunol. 2018 Apr 19;9:536.

30. Nirody JA, Budin I, Rangamani P. ATP synthase: Evolution, energetics, and membrane interactions. J Gen Physiol. 2020 Nov 2;152(11):e201912475.

31. Sohlenkamp C, Geiger O. Bacterial membrane lipids: diversity in structures and pathways. FEMS Microbiol Rev. 2016 Jan;40(1):133–59.

32. Mahammad S. & Parmryd I. (2015) in Methods in Membrane Lipids. Methods in Molecular Biology, ed. D. IO (Humana Press, New York, NY).

33. Horvath SE, Daum G. Lipids of mitochondria. Prog Lipid Res. 2013 Oct;52(4):590–614.

34. Mills EL, Kelly B, Logan A, Costa ASH, Varma M, Bryant CE, Tourlomousis P, Däbritz JHM, Gottlieb E, Latorre I, Corr SC, McManus G, Ryan D, Jacobs HT, Szibor M, Xavier RJ, Braun T, Frezza C, Murphy MP, O’Neill LA. Succinate Dehydrogenase Supports Metabolic Repurposing of Mitochondria to Drive Inflammatory Macrophages. Cell. 2016 Oct 6;167(2):457–470.e13.

35. Lou PH, Hansen BS, Olsen PH, Tullin S, Murphy MP, Brand MD. Mitochondrial uncouplers with an extraordinary dynamic range. Biochem J. 2007 Oct 1;407(1):129–40.

36. Kenwood B.M., Weaver J.L., Bajwa A., Poon I.K., Byrne F.L., Murrow B.A. et al. (2013) Identification of a novel mitochondrial uncoupler that does not depolarize the plasma membrane. Mol Metab. 3, 114–123.

37. Klose RJ, Kallin EM, Zhang Y. JmjC-domain-containing proteins and histone demethylation. Nat Rev Genet. 2006 Sep;7(9):715–27.

38. Liu PS, Wang H, Li X, Chao T, Teav T, Christen S, Di Conza G, Cheng WC, Chou CH, Vavakova M, Muret C, Debackere K, Mazzone M, Huang HD, Fendt SM, Ivanisevic J, Ho PC. α-ketoglutarate orchestrates macrophage activation through metabolic and epigenetic reprogramming. Nat Immunol. 2017 Sep;18(9):985–994.

39. Kruidenier L, Chung CW, Cheng Z, Liddle J, Che K, Joberty G, Bantscheff M, Bountra C, Bridges A, Diallo H, Eberhard D, Hutchinson S, Jones E, Katso R, Leveridge M, Mander PK, Mosley J, Ramirez-Molina C, Rowland P, Schofield CJ, Sheppard RJ, Smith JE, Swales C, Tanner R, Thomas P, Tumber A, Drewes G, Oppermann U, Patel DJ, Lee K, Wilson DM. A selective jumonji H3K27 demethylase inhibitor modulates the proinflammatory macrophage response. Nature. 2012 Aug 16;488(7411):404–8.

40. Becker L, Gharib SA, Irwin AD, Wijsman E, Vaisar T, Oram JF, Heinecke JW. A macrophage sterol-responsive network linked to atherogenesis. Cell Metab. 2010 Feb 3;11(2):125–35.

41. Ip WKE, Hoshi N, Shouval DS, Snapper S, Medzhitov R. Anti-inflammatory effect of IL-10 mediated by metabolic reprogramming of macrophages. Science. 2017 May 5;356(6337):513–519.

42. Dowling JK, Afzal R, Gearing LJ, Cervantes-Silva MP, Annett S, Davis GM, De Santi C, Assmann N, Dettmer K, Gough DJ, Bantug GR, Hamid FI, Nally FK, Duffy CP, Gorman AL, Liddicoat AM, Lavelle EC, Hess C, Oefner PJ, Finlay DK, Davey GP, Robson T, Curtis AM, Hertzog PJ, Williams BRG, McCoy CE. Mitochondrial arginase-2 is essential for IL-10 metabolic reprogramming of inflammatory macrophages. Nat Commun. 2021 Mar 5;12(1):1460.

43. Escoubet-Lozach L, Benner C, Kaikkonen MU, Lozach J, Heinz S, Spann NJ, Crotti A, Stender J, Ghisletti S, Reichart D, Cheng CS, Luna R, Ludka C, Sasik R, Garcia-Bassets I, Hoffmann A, Subramaniam S, Hardiman G, Rosenfeld MG, Glass CK. Mechanisms establishing TLR4-responsive activation states of inflammatory response genes. PLoS Genet. 2011 Dec;7(12):e1002401.

44. Brown MS, Goldstein JL. A proteolytic pathway that controls the cholesterol content of membranes, cells, and blood. Proc Natl Acad Sci U S A. 1999 Sep 28;96(20):11041–8.

45. Shen K, Pender CL, Bar-Ziv R, Zhang H, Wickham K, Willey E, Durieux J, Ahmad Q, Dillin A. Mitochondria as Cellular and Organismal Signaling Hubs. Annu Rev Cell Dev Biol. 2022 Oct 6;38:179–218.

46. Endo A, Kuroda M, Tanzawa K. Competitive inhibition of 3-hydroxy-3-methylglutaryl coenzyme A reductase by ML-236A and ML-236B fungal metabolites, having hypocholesterolemic activity. FEBS Lett. 1976 Dec 31;72(2):323–6.

## References

1. Benyoucef A. et al. UTX inhibition as selective epigenetic therapy against TAL1-driven T-cell acute lymphoblastic leukemia. Genes Dev. 30, 508–521 (2016).

2. Gutiérrez-Sanz O. et al. H2-Fueled ATP Synthesis on an Electrode: Mimicking Cellular Respiration. Angewandte Chemie, 6216–6220 (2016).

3. Mahammad S. & Parmryd I. Cholesterol depletion using methyl-β-cyclodextrin, in Methods in Membrane Lipids. Methods in Molecular Biology, Vol. 1232. (ed. I.O. D.) (Humana Press, New York, NY; 2015).

4. Bacia K., Scherfeld D., Kahya N. & Schwille P. Fluorescence correlation spectroscopy relates rafts in model and native membranes. Biophysical Journal 87, 1034–1043 (2004).

5. Chen P.S., Toribara J.T.Y. & H., W. Microdetermination of phosphorus. Analytical Chemistry, 1756-1758 (1956).

6. Amundson D.M. & Zhou M. Fluorometric method for the enzymatic determination of cholesterol. Journal of Biochemical and Biophysical Methods 38, 43–52 (1999).

7. Smith P.K. et al. Measurement of protein using bicinchoninic acid. Analytical Biochemistry 150, 76–85 (1985).

8. Pucadyil T.J. & Chattopadhyay A Cholesterol depletion induces dynamic confinement of the G-protein coupled serotonin1A receptor in the plasma membrane of living cells. Biochimica et Biophysica Acta (BBA) - Biomembranes 1768, 655–668 (2007).

9. Lanzetta P. A., Alvarez L. L. J., Reinach P S. & A., C.O. An improved assay for nanomole amounts of inorganic phosphate. Analytical Biochemistry 100, 95–97 (1979).

10. Kaim G. & Dimroth P. Construction, expression and characterization of a plasmid-encoded Na(+)-specific ATPase hybrid consisting of Propionigenium modestum FO-ATPase and Escherichia coli F1-ATPase. Eur J Biochem, 615–623 (1994).

11. GENCODE Consortium. (2021). GENCODE Mouse Release M25. Retrieved from https://www.gencodegenes.org/mouse/release_M25.html. Patro R, Duggal G, Love MI, Irizarry RA, Kingsford C (2017) Salmon provides fast and bias-aware quantification of transcript expression. Nat Methods 14:417.

12. Patro R, Duggal G, Love MI, Irizarry RA, Kingsford C (2017) Salmon provides fast and bias-aware quantification of transcript expression. Nat Methods 14:417.

13. Love MI, Huber W, Anders S. Moderated estimation of fold change and dispersion for RNA-seq data with DESeq2. Genome Biol. 2014;15(12):550.

14. Wu T, Hu E, Xu S, Chen M, Guo P, Dai Z, Feng T, Zhou L, Tang W, Zhan L, Fu x, Liu S, Bo X, Yu G (2021). “clusterProfiler 4.0: A universal enrichment tool for interpreting omics data.” The Innovation, 2(3), 100141.

15. Langmead B, Salzberg S. Fast gapped-read alignment with Bowtie 2. Nature Methods. 2012, 9:357–359.

16. Zhang Y, Liu T, Meyer CA, Eeckhoute J, Johnson DS, Bernstein BE, et al. Model-based Analysis of ChIP-Seq (MACS). Genome Biology. 2008;9(9):R137.

17. Stark R, Brown G (2011). DiffBind: differential binding analysis of ChIP-Seq peak data. http://bioconductor.org/packages/release/bioc/vignettes/DiffBind/inst/doc/DiffBind.pdf.

18. Ross-Innes CS, Stark R, Teschendorff AE, Holmes KA, Ali HR, Dunning MJ, Brown GD, Gojis O, Ellis IO, Green AR, Ali S, Chin SF, Palmieri C, Caldas C, Carroll JS. Differential oestrogen receptor binding is associated with clinical outcome in breast cancer. Nature. 2012 Jan 4;481(7381):389–93.

19. Love MI, Huber W, Anders S. Moderated estimation of fold change and dispersion for RNA-seq data with DESeq2. Genome Biol. 2014;15(12):550.

20. Robinson MD, McCarthy DJ, Smyth GK. EdgeR: a bioconductor package for differential expression analysis of digital gene expression data. Bioinformatics. 2010;26(1):139–40.

21. Wang Q, Li M, Wu T, Zhan L, Li L, Chen M, Xie W, Xie Z, Hu E, Xu S, Yu G (2022). “Exploring epigenomic datasets by ChIPseeker.” Current Protocols, 2(10), e585.

22. Yu G, Wang L, He Q (2015). “ChIPseeker: an R/Bioconductor package for ChIP peak annotation, comparison and visualization.” Bioinformatics, 31(14), 2382–2383.

23. Korotkevich G, Sukhov V, Sergushichev A (2019). “Fast gene set enrichment analysis.” bioRxiv. doi:10.1101/060012,

